# The tip of the VgrG spike is essential to functional type VI secretion assembly in *Acinetobacter baumannii*

**DOI:** 10.1101/808725

**Authors:** Juvenal Lopez, Pek Man Ly, Mario F. Feldman

**Affiliations:** Department of Molecular Microbiology, Washington University School of Medicine in St. Louis, St. Louis, MO, USA.

## Abstract

The type VI secretion system (T6SS) is a critical weapon in bacterial warfare between Gram-negative bacteria. Although invaluable for niche establishment, this machine represents an energetic burden to its host bacterium. *Acinetobacter baumannii* is an opportunistic pathogen that poses a serious threat to public health due to its high rates of multidrug resistance. In some *A. baumannii* strains, the T6SS is transcriptionally downregulated by large multidrug-resistance plasmids. Other strains, such as the clinical isolate AbCAN2, express T6SS-related genes but lack T6SS activity under laboratory conditions, despite not harboring these plasmids. This suggests that alternative mechanisms exist to repress the T6SS. Here, we employed a transposon mutagenesis approach in AbCAN2 to identify novel T6SS repressors. Our screen revealed that the T6SS of this strain is inhibited by a homolog of VgrG, an essential structural component of all T6SSs reported to date. We named this protein inhibitory VgrG (VgrGi). Biochemical and *in silico* analyses demonstrated that the unprecedented inhibitory capability of VgrGi is due to a single amino acid mutation in the widely conserved C-terminal domain of unknown function DUF2345. We also show that unlike in other bacteria, the C-terminus of VgrG is essential for functional T6SS assembly in *A. baumannii*. Our study provides insight into the architectural requirements underlying functional assembly of the T6SS of *A. baumannii*. We propose that T6SS-inactivating point mutations are beneficial to the host bacterium, as they eliminate the energy cost associated with maintaining a functional T6SS, which appears to be unnecessary for *A. baumannii* virulence.

**Importance:** Despite the clinical relevance of *A. baumannii,* little is known about its fundamental biology. Here, we show that a single amino acid mutation in VgrG, a critical T6SS structural protein, abrogates T6SS function. Given that this mutation was found in a clinical isolate, we propose that the T6SS of *A. baumannii* is likely not involved in virulence, an idea supported by multiple genomic analyses showing that the majority of clinical *A. baumannii* strains lack proteins essential to the T6SS. We also show that, unlike in other species, the C-terminus of VgrG is a unique architectural requirement for functional T6SS assembly in *A. baumannii,* suggesting that over evolutionary time, bacteria have developed changes to their T6SS architecture, leading to specialized systems.

## Introduction

Bacterial life involves constant interactions between diverse bacterial species. These bacteria-bacteria interactions underlie the formation of complex bacterial communities, which are the subject of constant compositional changes due to factors, such as nutrient availability and interbacterial antagonism (1, 2). Several seminal studies have shown that it is essential for bacterial pathogens to outcompete the normal host microbiota to establish a niche and cause disease (3–5). Interbacterial competition has also been shown to facilitate the exchange of genetic material, thus promoting the dissemination of antibiotic resistance genes and virulence factors (6–9). Understanding how bacteria compete with one another provides insight into a critical aspect of bacterial life.

The type VI secretion system (T6SS) mediates interbacterial competition between Gram-negative bacteria (2, 10, 11). It is composed of a minimum of thirteen conserved structural proteins (TssA-TssM), all encoded in a single locus (12). These constituents assemble into a trans-envelope, bacteriophage tail-like structure that delivers toxic effector proteins into adjacent bacterial cells (13, 14). The “tail” is a cytosolic complex composed of a tube of Hcp (or TssD) hexamers topped with a spike of three VgrG (or TssI) proteins and, in some cases, a single PAAR protein. Hcp, VgrG and PAAR bind antibacterial T6SS effectors, which target essential cell structures, such as the cell wall, genetic material or cell membrane (15, 16). The spiked tube is in turn encompassed by a contractile sheath comprised of TssB and TssC. When the sheath contracts, it propels the effector-loaded spiked tube outward from the attacking cell and into adjacent cells. Following its contraction, the sheath is disassembled by the ATPase ClpV (or TssH) to enable sheath constituents to be reused in subsequent T6SS assemblies (17).

The dynamic T6SS tail is built from a multimeric protein complex known as the baseplate and is anchored to the bacterial envelope by a membrane complex (18, 19). The membrane complex is composed of three membrane-associated proteins: TssJ, TssM and TssL (19). In contrast, the baseplate forms in the cytoplasm, beginning with the trimerization of VgrG and the subsequent assembly of wedge proteins (TssE-G and TssK) around the VgrG hub (20–22). Following its assembly, the baseplate interacts with the membrane complex (18, 22, 23) and the tube and sheath assemble at the N-terminus of VgrG in a highly coordinated process facilitated by TssA (18, 24).

The T6SS is energetically costly (26). Thus, many T6SS-encoding bacteria tightly regulate their T6SS at the transcriptional, posttranscriptional and/or posttranslational levels (27, 28). The complex regulatory networks governing T6SS activity enable bacteria to respond according to environmental signals, including: temperature, pH, cation/nutrient availability, osmolarity and membrane perturbation (27, 29–37). For instance, the antibacterial T6SS of *Pseudomonas aeruginosa* (HSI-I), regarded as a defensive T6SS, remains silenced unless *P. aeruginosa* perceives an attack from a competing bacterium (38). When an attack is perceived, a posttranslational threonine phosphorylation pathway activates the forkhead-associated domain-containing protein Fha1, ultimately resulting in the assembly of a functional T6SS in the location of the perceived attack (38). Thus, *P. aeruginosa* overcomes the high energy cost of the T6SS by preventing futile T6SS attacks in the absence of competing bacteria. However, the mechanisms underlying T6SS regulation in other less-studied medically relevant bacteria remain poorly understood.

*Acinetobacter baumannii* is a Gram-negative bacterial pathogen with alarming rates of multidrug resistance and mortality (39). We have previously shown that *A. baummanii* strains encode a single, highly conserved T6SS locus (40). Despite the notable genetic conservation, diverse *A. baummanii* strains possess varying levels of Hcp secretion, the hallmark of T6SS activity. One subset of strains secretes Hcp under standard laboratory conditions, indicating that they possess a constitutively active (or offensive) T6SS. A second subset of strains possess a silenced T6SS because they harbor a family of multidrug-resistance plasmids named Large Conjugative Plasmids (LCPs), which transcriptionally repress the T6SS locus (41–43). The final subset of *A. baumannii* strains expresses T6SS-related genes and does not harbor an LCP but does not secrete Hcp under standard laboratory conditions (40, 44–46). These strains are expected to regulate their T6SS by as-yet uncharacterized mechanisms. The clinical *A. baummanii* strain CAN2 (AbCAN2, formerly named Ab1225) expresses Hcp but does not secrete it under standard laboratory conditions (40). Here, we report the whole-genome sequence of AbCAN2 and determine that it encodes all genes necessary for a functional T6SS and lacks an LCP. Using a transposon mutagenesis approach, we identified a VgrG homolog as an unexpected inhibitor of the T6SS of AbCAN2. Our bioinformatics and biochemical analyses provide insight into the characteristics that differentiate an inhibitory VgrG from a canonical VgrG. Furthermore, we demonstrate a previously unappreciated role of the C-terminus of VgrG proteins in T6SS assembly in *A. baummanii*.

## Results

### Clinical strain AbCAN2 encodes a T6SS locus

AbCAN2 is a coccygeal isolate collected at the University of Alberta Hospital, Canada with an inactive T6SS under standard laboratory conditions (40). To gain a better understanding of why the T6SS is inactive in this strain, we determined the whole-genome sequence of AbCAN2 (BioSample accession: SAMN12912378). We found that this strain does not harbor an LCP. Additionally, it encodes all genes required for a functional T6SS. In fact, the T6SS locus of AbCAN2 is syntenic with that of other *A. baumannii* strains, including the lab strain *A. baumannii* ATCC 17978 (Ab17978) (Fig. 1) (40). The T6SS genes of AbCAN2 and Ab17978 present remarkably high levels of sequence conservation, ranging from 98% to 100% nucleotide identity (Table S1). Besides encoding 12 of the 13 structural T6SS proteins conserved among Proteobacteria (*Acinetobacter* species lack a TssJ homolog (40)), the T6SS locus of AbCAN2 also contains genes coding for accessory proteins TagF, TagN, PAAR and TagX. Additionally, the T6SS locus harbors genes that code for three hypothetical proteins that are conserved among *Acinetobacter* species, some of which have been shown to be essential for T6SS function (47, 48). The specific roles of these proteins are yet to be elucidated.

**Figure 1.**
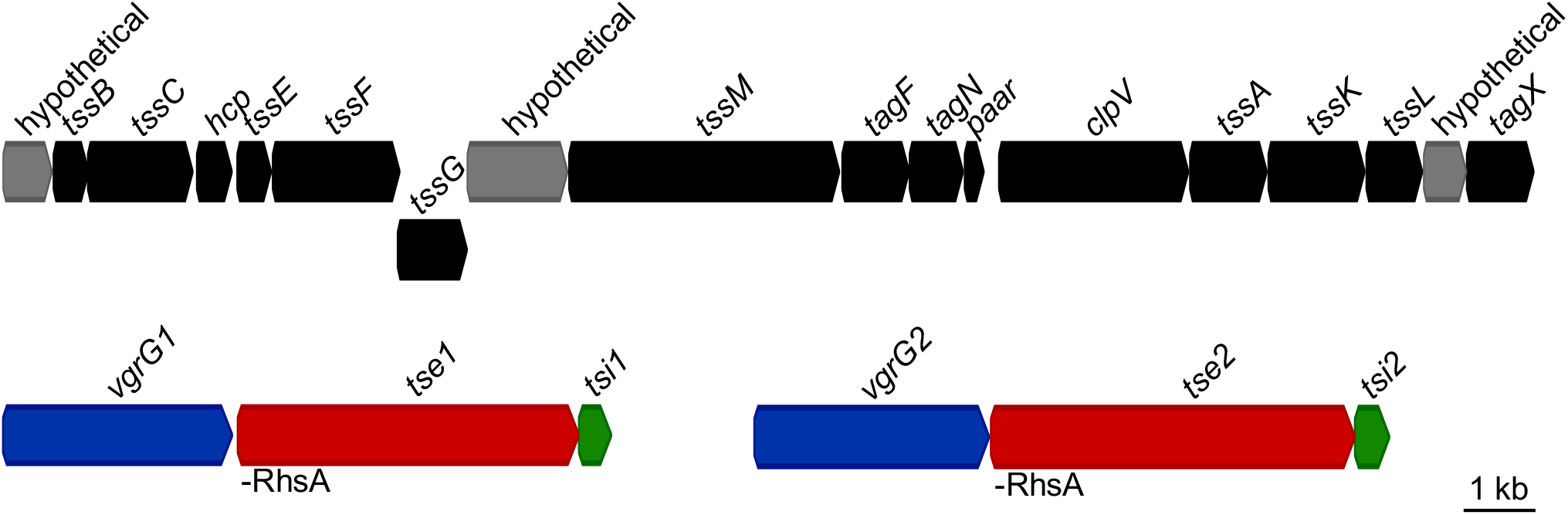
AbCAN2 encodes conserved T6SS-related genes. Schematic of the T6SS locus and *vgrG* gene clusters encoded by AbCAN2. Functional domains identified in *tse1* and *tse2* are shown.

AbCAN2 also encodes 2 *vgrG* gene clusters located remotely from its T6SS locus (Fig. 1). We arbitrarily named the VgrGs of each cluster VgrG1 and VgrG2. Each VgrG is encoded upstream of an open reading frame (ORF) containing a predicted rearrangement hotspot A (RhsA) domain. T6SS effectors of the RhsA family have been previously reported to possess DNase activity (49–51); thus, it is possible that the proteins coded by these ORFs possess nuclease activity. Given their genetic proximity to *vgrG1* and *vgrG2*, we expect these proteins to be T6SS effectors. Thus, we named these ORFs type six effector 1 (*tse1*) and *tse2*, respectively. Downstream of each *tse1* and *tse2* is an ORF that we predict to be the cognate immunity proteins of these effectors. Immunity proteins are commonly encoded downstream of their cognate effector, and they inactive their cognate effector by specifically binding it and obstructing its active site (10). Thus, we named these ORFs type six immunity 1 (*tsi1*) and *tsi2*, respectively.

### A VgrG homolog inhibits the T6SS of AbCAN2

Our finding that AbCAN2 lacks T6SS activity despite encoding all genes required for T6SS assembly and not harboring an LCP led us to hypothesize that AbCAN2 encodes a novel T6SS repressor. To identify the putative novel repressor of the T6SS, we generated a transposon mutant library and screened mutants for Hcp secretion, using our high-throughput Hcp colony blot assay (see Materials and Methods) (52). We hypothesized that genetic disruption of a T6SS inhibitor would result in appreciable T6SS activity in a mutant strain. Of the ∼3000 colonies screened, we detected Hcp in culture supernatants of four unique mutant strains, indicating that these transposon mutants possess an active T6SS (Fig. 2a). Surprisingly, the gene disrupted in all four mutant strains was *vgrG1* (Fig. 2b). VgrG is a conserved structural component of all T6SSs reported to date. It is a modular protein containing N-terminal domains structurally similar to spike complex proteins of bacteriophage contractile tails (gp27 and gp5) as well as C-terminal DUF2345 and transthyretin (TT)-like domains, whose roles are yet to be fully understood (20, 21, 53–55). VgrGs are widely regarded as essential to T6SS assembly and function, being implicated in roles such as baseplate formation, proper Hcp tube assembly and effector delivery (18, 22, 24, 56). Thus, our result that a VgrG homolog inhibits the T6SS is against the current paradigm of VgrG function. To confirm our results, we generated a clean deletion mutant of *vgrG1* (hereafter referred to as *vgrGi* for inhibitory VgrG) and compared its T6SS activity to that of wild-type (WT) AbCAN2, using Hcp secretion and bacterial killing assays. Consistent with our previous result, Hcp was detected in the supernatant fraction of *ΔvgrGi* but not WT AbCAN2 or the complemented strain (*vgrGi+*), indicating that VgrGi inhibits the T6SS of AbCAN2 (Fig. 2c, Fig. S1a). Similarly, AbCAN2*ΔvgrGi*, but not WT or *vgrGi+*, demonstrated significant levels of interbacterial killing when co-incubated with *E. coli* (Fig. 2d, Fig. S1b). Altogether, our data indicate that VgrGi inhibits T6SS activity in AbCAN2.

**Figure 2.**
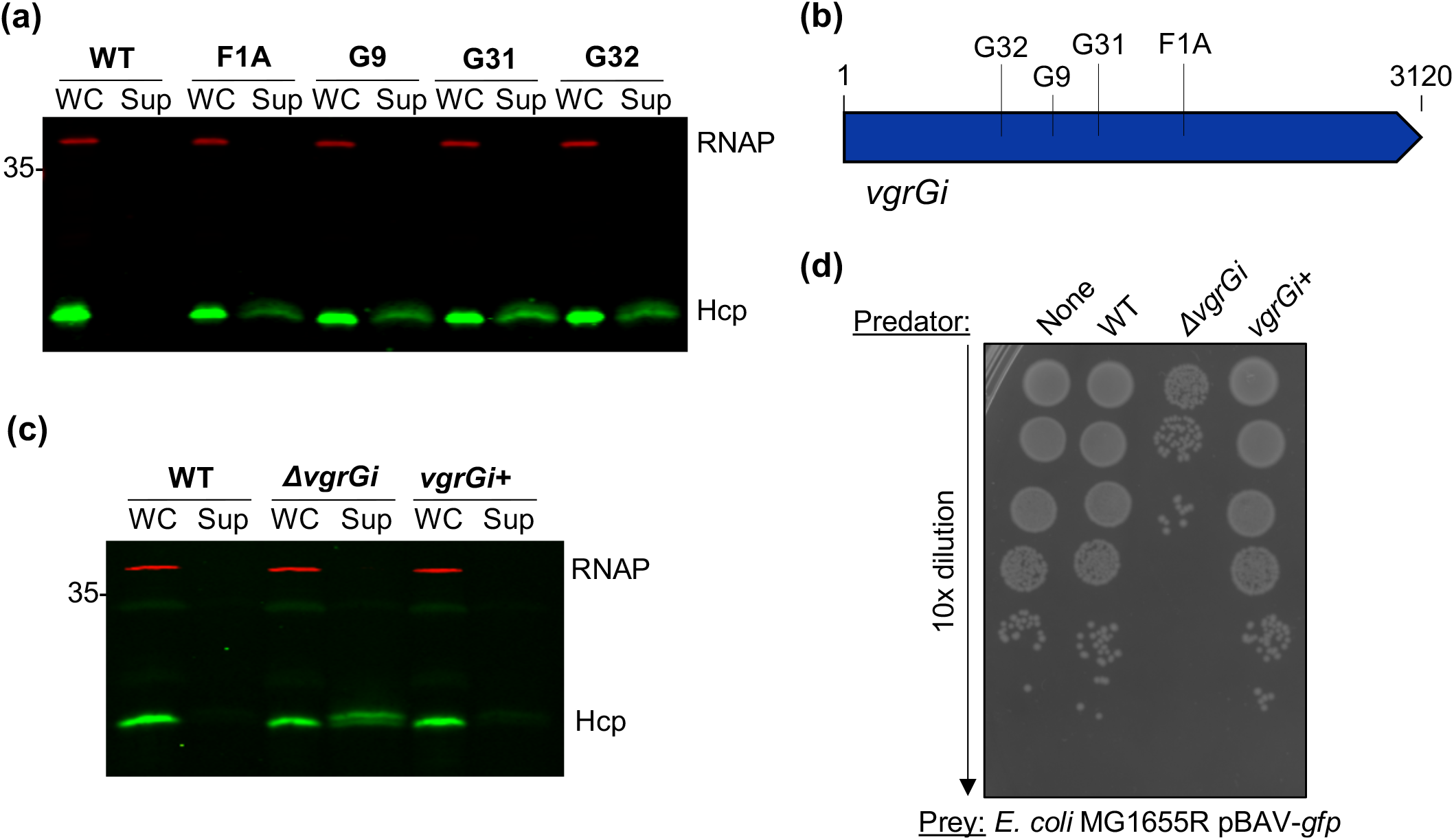
VgrGi inhibits the T6SS of AbCAN2. **a**, Western blot of OD-normalized whole cell (WC) and supernatant (Sup) fractions of wild-type (WT) or unique transposon insertion mutants of AbCAN2 probing for Hcp expression and secretion. **b**, Schematic of transposon insertion sites within *vgrGi.* G32: A853, G9: A1108, G31: A1362, F1A: A1846. **c**, Western blot probing for Hcp expression and secretion in AbCAN2 WT, *ΔvgrGi*, or the complemented strain (*vgrGi+*). **a**, **c**, RNAP is included as a lysis and loading control. **d**, Representative T6SS killing assay. Serial dilutions of *E. coli* were spotted after a 3.5-h incubation with the indicated predator strains at a 10:1 (predator:prey) ratio. Statistical analysis is shown in Fig. S1b. **c**, **d**, WT and *ΔvgrGi* harbor empty vector pWH1266.

### VgrGi acts independently to inhibit the T6SS

A previous report in *Vibrio cholerae* suggests that T6SS effectors may play a critical role in T6SS assembly (57). Thus, we tested whether AbCAN2 effectors Tse1 and Tse2 or VgrG2 were involved in T6SS inhibition. To this end, we generated mutant strains of AbCAN2 lacking *tse/tsi1*, *tse/tsi2*, *vgrG2* or both *vgrGi* and *vgrG2* (*ΔvgrGi*,*2*) and determined their T6SS activity by Hcp secretion assay. We did not detect Hcp in the supernatant fraction of any of the mutant strains, indicating that VgrGi-mediated T6SS inhibition is independent of any proteins encoded within the two *vgrG* clusters of AbCAN2 (Fig. 3). Importantly, we also found that unlike *ΔvgrGi*, the *ΔvgrGi*,*2* double mutant does not secrete Hcp. This result demonstrates that VgrG2 acts as a canonical VgrG; it is required for proper T6SS function and likely participates in the assembly of the machine.

**Figure 3.**
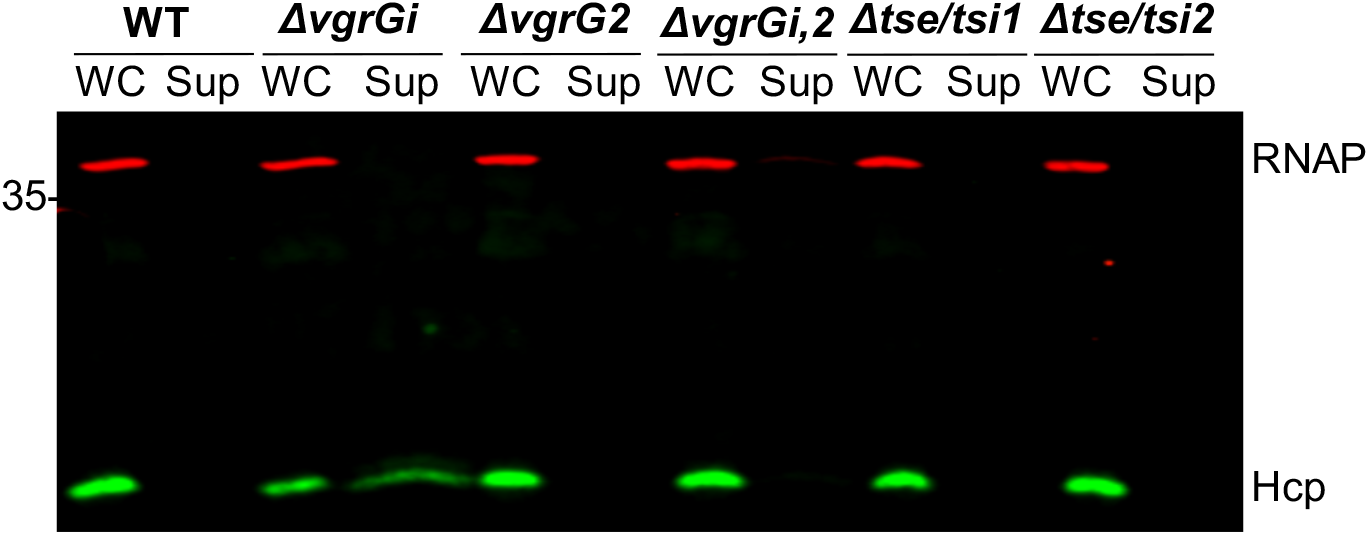
Additional VgrG-related proteins are not involved in T6SS inhibition. Western blot of OD-normalized whole cell (WC) and supernatant (Sup) fractions of wild-type (WT) AbCAN2 or the indicated mutants probing for Hcp expression and secretion. RNAP is included as a lysis and loading control.

Next, we tested whether VgrGi-mediated T6SS inhibition is due to an intrinsic property of AbCAN2 or whether VgrGi can also inhibit the T6SS of a heterologous host. To this end, we expressed plasmid-borne 6xHis-tagged VgrGi in Ab17978 lacking LCP pAB3 (hereafter referred to as WT Ab17978), which possesses constitutive T6SS activity under standard laboratory conditions (41), and probed for Hcp secretion. We found that WT Ab17978 expressing VgrGi secreted similar levels of Hcp compared with the vector control (Fig. 4a, Fig. S2). This result suggested that VgrGi is unable to inhibit the T6SS of a non-cognate host. Nonetheless, it is noteworthy that unlike AbCAN2, which encodes one canonical VgrG, VgrG2, Ab17978 encodes four VgrG homologs, each sufficient for assembling a functional T6SS (47). We reasoned that it was possible that VgrGi was unable to inhibit the T6SS of Ab17978 due to functional redundancy between the four VgrGs. Therefore, we then expressed VgrGi in mutant strains of Ab17978 lacking one to three VgrGs. We found that VgrGi expression in Ab17978*ΔvgrG1* resulted in a considerable reduction of Hcp detected in culture supernatants (Fig. 4b). Remarkably, VgrGi completely abolished Hcp secretion in Ab17978 *ΔvgrG1,2* and *ΔvgrG1,2,3* (Fig. 4c-d), indicating that the inhibitory capability of VgrGi extends beyond AbCAN2.

**Figure 4.**
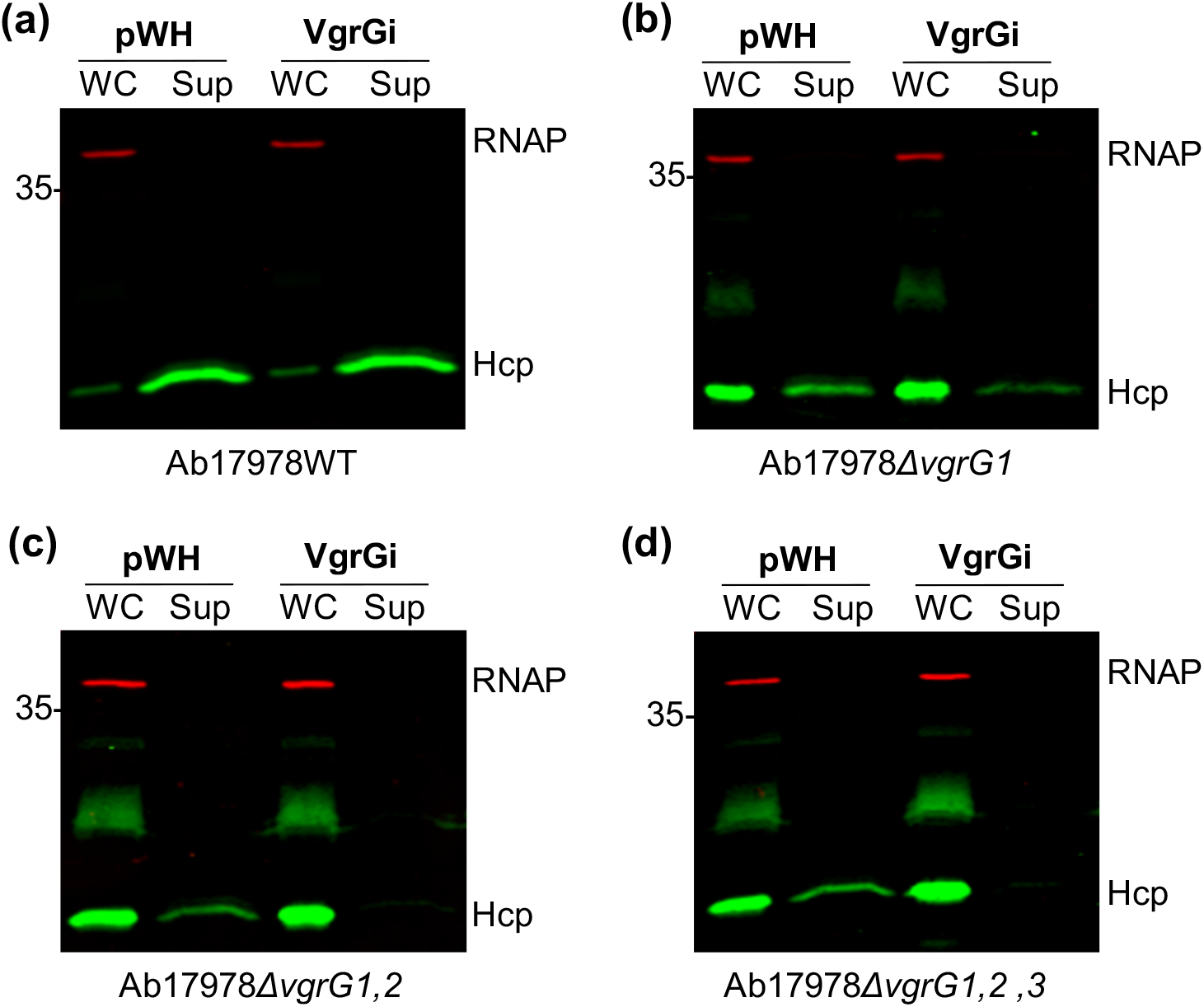
VgrGi inhibits the T6SS of VgrG mutant strains of Ab17978. Western blot of OD-normalized whole cell (WC) and supernatant (Sup) fractions of wild-type (WT) Ab17978 or the indicated *vgrG* mutant strains probing for Hcp expression and secretion. RNAP is included as a lysis and loading control.

Taken together, our results show that VgrGi can inhibit the T6SS of its native strain as well as that of Ab17978 mutant strains with limited VgrGs. Given that AbCAN2 and Ab17978, possess a different arsenal of effectors, these findings provide further evidence that VgrGi-mediated T6SS inhibition is independent of effectors expressed.

### VgrGi-mediated T6SS inhibition is due to a leucine-to-arginine mutation

VgrGi is, to the best of our knowledge, the first VgrG homolog with an ability to inhibit the T6SS. Thus, we sought to identify unique characteristics that differentiate VgrGi from other VgrGs. However, protein sequence and structural prediction analyses revealed that VgrGi contains domains similar to those found in canonical VgrGs of diverse bacteria, such as VgrG1 of *P. aeruginosa* PAO1 (VgrG1^Pa^) and VgrG1 of EAEC 17-2 (VgrG1^Ec^) (53, 55, 58). (Despite their identical name, VgrG1^Pa^ and VgrG1^Ec^ are two distinct proteins.) VgrGi contains two domains at its N-terminus that closely resemble bacteriophage T4 tail spike proteins gp27 (residues 1-426) and gp5 (residues 537-666) (Fig. 5a). These domains are linked by an oligosaccharide-binding (OB)-fold domain (residues 427-536). The C-terminus of VgrGi is composed of a DUF2345 domain (residues 688-834) followed by a TT-like domain (residues 860-907) and a region of ∼130 residues with no predicted functional domains. Finally, the N- and C-terminal segments of VgrGi are linked by a predicted coiled-coil region (residues 667-687) (Fig. 5a).

**Figure 5.**
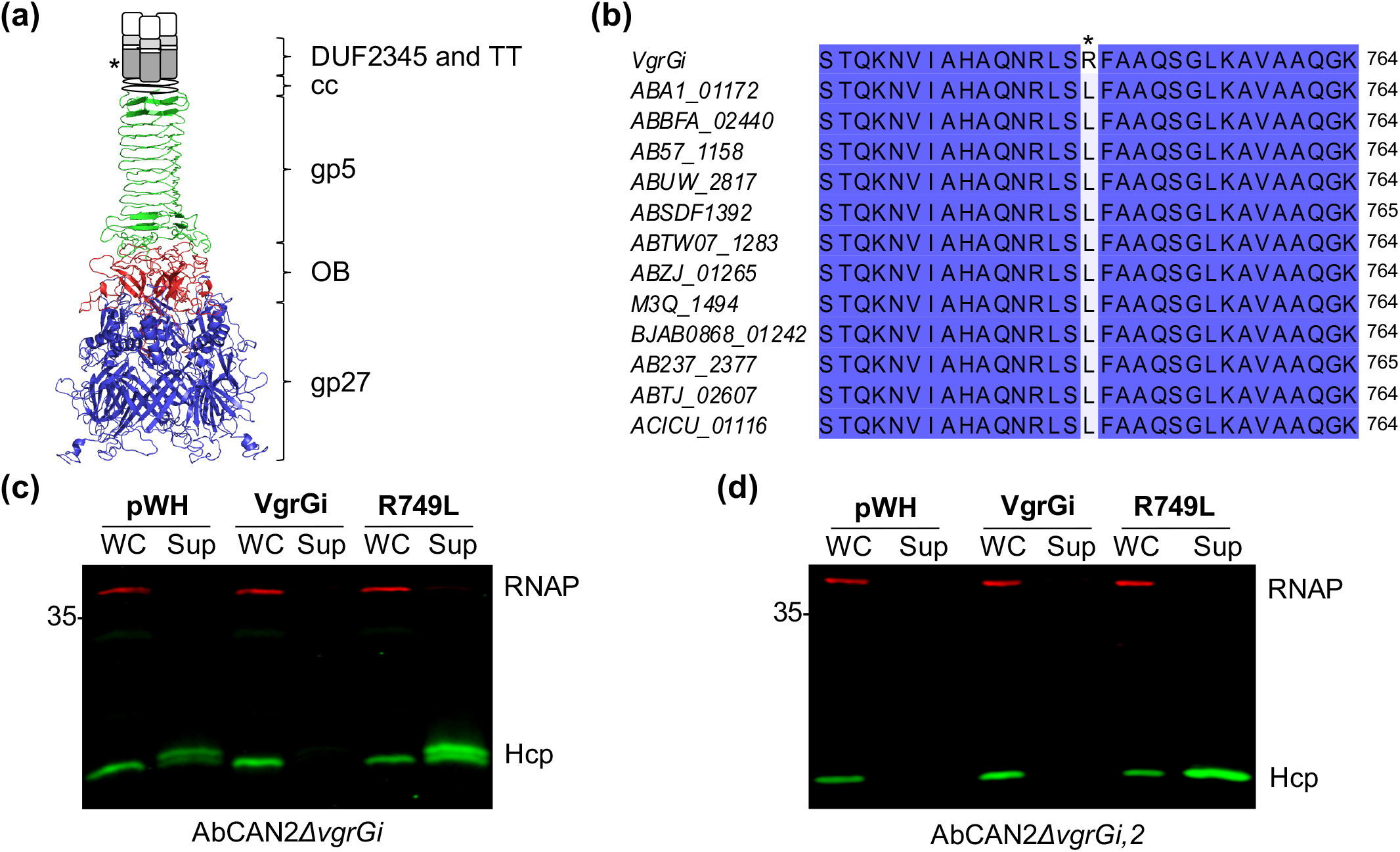
Arginine residue in DUF2345 domain of VgrGi is necessary for T6SS inhibition. **a**, Schematic of the predicted structure of VgrGi based on the crystal structure of the *P. aeruginosa* VgrG1 (PDB: 4MTK). Different domains are represented in different colors: gp27 (blue), OB-fold (red) and gp5 (green). The DUF2345 (dark gray) and transthyretin (TT, light gray) could not be modelled and are represented by rectangles. Patterned and white rectangles represent areas for which no domain is predicted. The conceptualization of this figure borrowed heavily from M. G. Renault *et al.,* J Mol Biol 430:3143-3156, 2018, https://doi.org/10.1016/j.jmb.2018.07.018. **b**, Amino acid alignment of VgrGi with other Class 1 VgrGs, according to the nomenclature of Fitzsimmons *et al.*, Infect Immun 86:e00297-18, 2018, https://doi.org/10.1128/IAI.00297-18. Identical residues are highlighted in blue. **a, b**, The asterisk highlights the L749R mutation present in VgrGi relative to other VgrGs of the same phylogenetic group. **c**, **d**, Western blot probing for Hcp expression and secretion. Shown are OD-normalized whole cell (WC) and supernatant (Sup) fractions of AbCAN2*ΔvgrGi* (**c**) or *ΔvgrGi,2* (**d**) expressing VgrGi, VgrGi R749L or a vector control (pWH). RNAP is included as a lysis and loading control.

Given that we identified no differences in the predicted functional domains of canonical VgrGs and those of VgrGi, we reasoned that VgrGi may possess critical differences in its primary sequence that are responsible for its inhibitory ability. VgrGs encoded by *A. baumannii* strains were recently grouped into six distinct phylogenetic groups (59). Based on amino acid identity (>95%), we determined that VgrGi belongs to Class 1 VgrGs. Interestingly, despite such remarkable levels of sequence conservation, we found that VgrGi contains a leucine-to-arginine substitution in position 749 compared with all other Class 1 VgrGs (Fig. 5b). We hypothesized that the leucine-to-arginine mutation was responsible for the unprecedented ability of VgrGi to inhibit the T6SS of *A. baumannii*. To test our hypothesis, we expressed plasmid-borne 6xHis-tagged VgrGi or a VgrGi revertant mutant containing an arginine-to-leucine mutation (hereafter referred to simply as R749L) in AbCAN2*ΔvgrGi* and probed for Hcp secretion. As expected, we detected Hcp in the supernatant of the vector control but not in the *vgrGi+* strain (Fig. 5c). Remarkably, *ΔvgrGi* expressing R749L (*vgrGi*_R749L_*+*) retained T6SS activity (Fig. 5c, Fig. S3a), indicating that the mutation R749L abrogated the inhibitory capability of VgrGi. In fact, *vgrGi*_R749L_*+* showed higher levels of Hcp secretion compared with the vector control (Fig. 5c), suggesting that R749L likely participates in T6SS assembly. To test this, we expressed plasmid-borne 6xHis-tagged VgrGi or R749L in AbCAN2*ΔvgrGi,2*. This strain lacks T6SS activity (Fig. 3) but encodes all other genes necessary to assemble a functional T6SS. Thus, we hypothesized that if R749L participates in T6SS assembly, heterologous expression of this protein in *ΔvgrGi,2* would result in T6SS activity. Indeed, *ΔvgrGi,2* expressing R749L but not VgrGi or empty vector secretes Hcp (Fig. 5d, Fig. S3b), indicating that the mutation R749L converts VgrGi to a canonical VgrG essential to T6SS assembly.

Altogether, our results indicate that VgrGi is lacks any functional domains that differentiate it from canonical VgrGs. Instead, its inhibitory capability is due to a single amino acid mutation.

### Polar or charged residues within the SLFAAQ motif disrupt VgrG function

Our previous result suggests that a leucine residue in position 749 plays an important role in canonical VgrG function. A protein alignment analysis revealed that despite considerable levels of divergence among VgrGs of diverse *A. baumannii* strains, most VgrGs possess a conserved SLFAAQ motif (Fig. S4). The conservation of this motif further points to the leucine residue as important for VgrG function. Thus, we hypothesized that substitution of the conserved leucine for arginine would convert a canonical VgrG to an inhibitory VgrG. VgrG1 of Ab17978 (ACX60_17665) possesses a leucine residue within its SLFAAQ motif, L758, and is important for T6SS function (Fig. S4) (47). To test our hypothesis, we generated plasmid-borne 6xHis-tagged VgrG1 (*vgrG1*+) or VgrG1 bearing a L758R mutation (*vgrG1*_L758R_+), expressed them in Ab17978*ΔvgrG1* (Fig. S5) and determined the resulting levels of Hcp secretion. In contrast with the *vgrG1*+ strain, which had increased levels of Hcp secretion compared with the vector control, *vgrG1*_L758R_+ demonstrated a marginal level of Hcp secretion (Fig. 6). This result indicates that substitution of a conserved leucine residue for arginine confers VgrG1 the ability to repress the T6SS of Ab17978.

**Figure 6.**
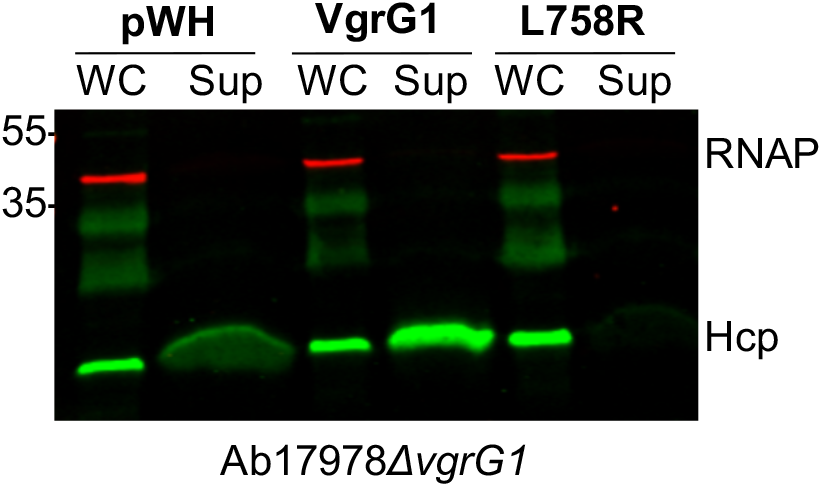
Leucine-to-arginine substitution converts canonical VgrG1 to an inhibitory VgrG. Western blot of Ab17978*ΔvgrG1* expressing pWH (empty vector), VgrG1 or VgrG1 L758R probing for Hcp expression and secretion. To facilitate visualization of Hcp in the supernatant fractions pWH and L758R, twice the amount of supernatant (normalized by OD) from these strains were loaded relative to Ab17978*ΔvgrG1* expressing VgrG1. RNAP is included as a lysis and loading control.

Next, we tested whether VgrGi-mediated T6SS inhibition depends on a specific property of arginine or whether other amino acids yield similar phenotypes. To this end, we generated pWH1266 vector constructs containing the following point mutants of VgrGi: R749K, R749D, R749N and R749F. Then, we expressed these mutants in AbCAN2*ΔvgrGi* and determined whether expression of these constructs results in T6SS inhibition (Fig. S6). We found that VgrGi variants containing polar or charged residues (i.e. R749K, R749D and R749D) inhibit Hcp secretion (Fig. 7). In contrast, VgrGi variants containing nonpolar residues (i.e. R749L and R749F) do not inhibit Hcp secretion (Fig. 7). Similar results were obtained when these constructs were expressed in Ab17978*ΔvgrG1,2* (Fig. S7). Collectively, our results indicate that polar or charged residues in the second position of the SLFAAQ motif disrupt the function of VgrGs in T6SS assembly. In fact, mutants with these inactivating mutations have a dominant negative effect on T6SS function.

**Figure 7.**
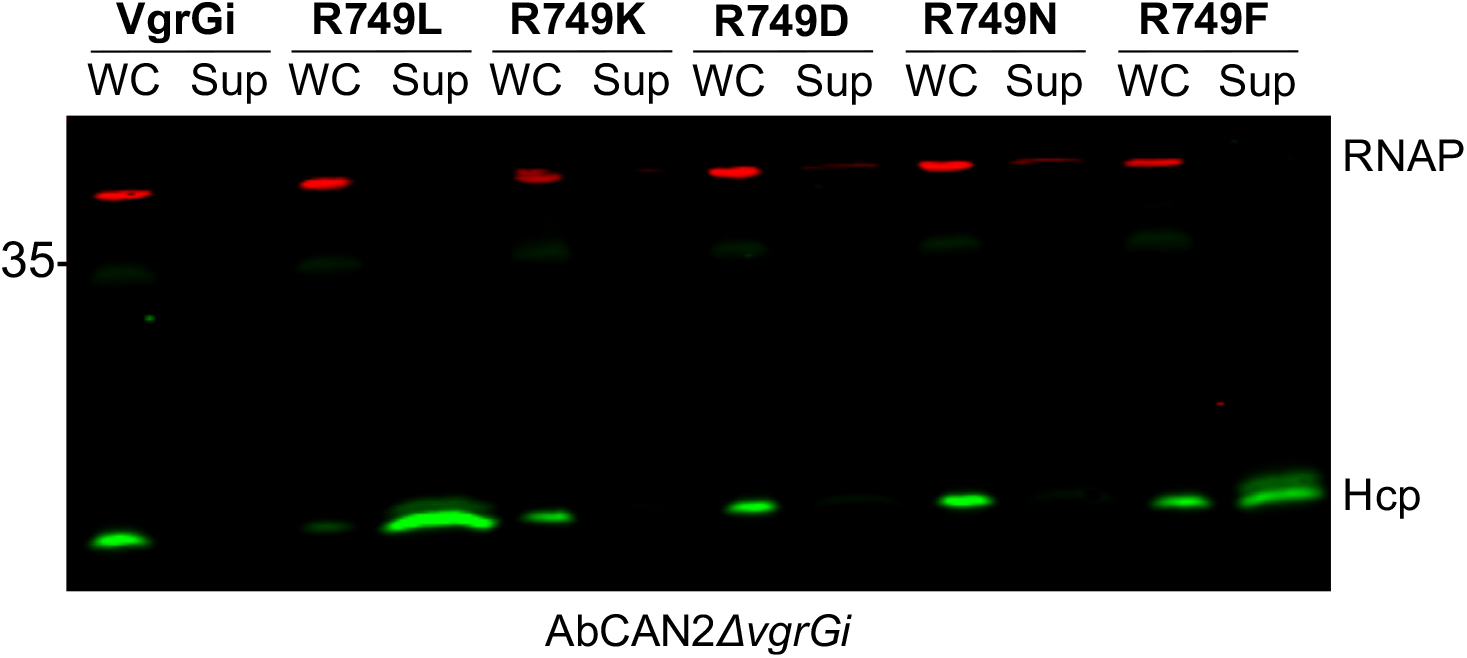
Polar and charged residues in position 749 result in T6SS inhibition by VgrGi. Western blot probing for Hcp expression and secretion. Shown are OD-normalized whole cell (WC) and supernatant (Sup) fractions of AbCAN2*ΔvgrGi* expressing the indicated VgrGi point mutants. RNAP is included as a lysis and loading control.

### The C-terminus of VgrG is essential to T6SS function in *A. baumannii*

The SLFAAQ motif lies within the C-terminal DUF2345 domain of VgrGi. Although present in VgrGs of diverse Gram-negative bacteria, the role of the DUF2345 domain in T6SS assembly remains poorly understood. Recent work in *E. coli* showed that the gp27 domain of VgrG1^Ec^ is sufficient for T6SS activity, indicating that the DUF2345 and TT domains are dispensable for T6SS assembly (53). Similarly, the C-terminus of *P. aeruginosa* VgrG5 is not required to support VgrG4b secretion (54). Nonetheless, our results indicate that a point mutation within the C-terminal DUF2345 domain of *A. baumannii* VgrGs is sufficient to disrupt T6SS activity. Thus, we hypothesized that in *A. baumannii*, the DUF2345 domain plays an essential role in the assembly of a functional T6SS. To test our hypothesis, we expressed different truncations of R749L in AbCAN2*ΔvgrGi,2* and determined the resulting levels of Hcp secretion. As shown in Figure 5d, *ΔvgrGi,2* lacks T6SS activity unless it expresses a functional VgrG (e.g. full length R749L). The R749L constructs tested include: the gp27 domain (gp27), a C-terminal truncation ending with the gp5 domain (C-gp5), a C-terminal truncation ending with the DUF2345 domain (C-DUF_R749L_) and mutants lacking either the DUF2345 domain (ΔDUF) or the TT domain (ΔTT_R749L_) but possessing all other domains (Fig. 8a, Fig. S8). We found that in contrast to full length R749L, expression of none of the R749L truncations tested led to Hcp secretion (Fig. 8b). This result indicates that the C-terminus of R749L is essential to the role of VgrG in assembling a functional T6SS.

**Figure 8.**
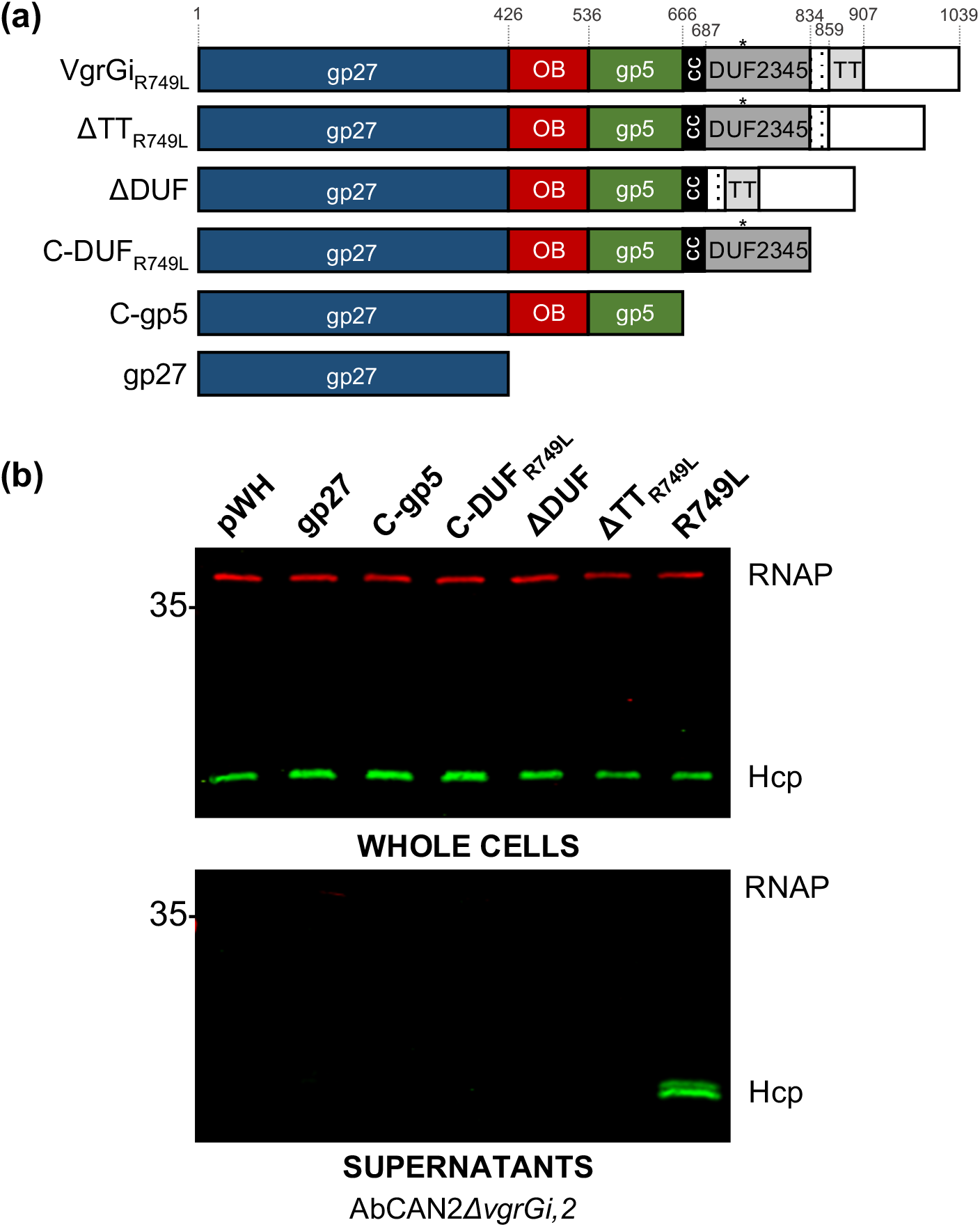
C-terminus of VgrGi_R749L_ is required for T6SS activity. **a**, Schematic of the domains present in VgrGi_R749L_ as well as the truncations made for the experiment in b. Amino acid boundaries are indicated and the color scheme is consistent with that of Fig. 5a. The asterisk indicates the position of the R749L mutation. **b**, Western blot probing for Hcp expression (top) and secretion (bottom) in AbCAN2*ΔvgrGi* expressing the indicated VgrGi_R749L_ truncations. RNAP is included as a lysis and loading control.

## Discussion

The T6SS is a critical weapon for interbacterial warfare (12). Seminal studies have provided general principles that are largely applicable to the T6SS of various bacterial species. However, it is becoming increasingly clear that the T6SS is a diversified machine, with important structural and functional differences depending on the bacterial host (32, 60, 61). Here, we identified a VgrG homolog with an unprecedented ability to inhibit the T6SS of the clinical isolate AbCAN2. Our efforts to characterize the mechanism underlying VgrGi-mediated T6SS inhibition led us to determine that the C-terminus of VgrG is essential for functional T6SS assembly in *A. baumannii*.

VgrG is one of the most versatile proteins of the T6SS. It is an essential structural component of the baseplate and sharpens the Hcp tube to facilitate penetration into adjacent cells and delivery of effectors (21, 22, 55). Despite its relevance, VgrG remains largely understudied compared with T6SS proteins likely due to its inherent insolubility (62). Only one full length VgrG protein has been structurally characterized, VgrG1^Pa^ (58). Notably, VgrG1^Pa^ is much shorter compared with other VgrGs (643 aa versus 841 aa of VgrG1^Ec^ and 1039 aa of VgrGi). Thus, the functional domains present in longer VgrGs remain largely uncharacterized.

One of the functional domains present in longer VgrGs is the DUF2345 domain. In *E. coli*, this domain has been shown to stabilize the interaction between the TT domain of VgrG1 and effector Tle1, but it is dispensable for T6SS assembly and dynamics (53, 55). Despite its seeming dispensability for T6SS assembly, this domain is present in VgrGs of a wide range of Gram-negative bacteria, including: *V. cholerae* (VCV52_2925), *Klebsiella pneumoniae* (BN49_3373), *P. aeruginosa* (PA0262) and *Salmonella enterica* (FJR63_23580). In fact, VgrGs were originally described as possessing a COG4253 functional domain, which belongs to the DUF2345 superfamily (61). Our results suggest that the DUF2345 domain may play a role in T6SS assembly in *A. baumannii*. However, as C-DUF_R749L_ is not sufficient for T6SS assembly (Fig. 8), the DUF2345 domain may require additional elements present at the C-terminus of VgrG for proper function. Moreover, a recent bioinformatic analysis of 73 VgrGs across 22 *A. baumannii* strains as well as the soil bacterium *A. baylyi* ADP1 showed that all VgrGs contain a DUF2345 domain (59). The high levels of conservation suggest that the DUF2345 domain may play a particularly important role in the T6SS of *Acinetobacter* species.

It is noteworthy that our results are not in conflict with previous results indicating that the C-terminus of VgrG is dispensable for T6SS function (53, 54). Instead, we propose that over evolutionary time, bacteria developed changes to their T6SS architecture, leading to specialized systems. Indeed, it was recently reported that TssA homologs carry out three diverse functions depending on their C-terminal domain (63). Moreover, homologs of membrane complex protein TssM possess a Walker A motif; however, no general role for this motif has been described. In *Agrobacterium tumefaciens*, for instance, TssM exhibits ATP binding and hydrolysis as well as ATP-binding-dependent conformational changes, all of which are implicated in T6SS function (64, 65). In contrast, the Walker A motif of the *Edwardsiella tarda* TssM, is dispensable for T6SS activity (66). Thus, it seems likely that bacteria have co-opted conserved structural proteins to better suit their needs.

Future work will characterize the mechanism by which VgrGi inhibits the T6SS, as it may provide insight into the role of the C-terminus of *A. baumannii* VgrGs. Our finding that VgrGi inhibits the T6SS of Ab17978 only when endogenous VgrGs are limited suggests that VgrGi acts at the post-translational level. We propose that VgrGi likely inhibits the T6SS by preventing its assembly. VgrGs can form heterotrimeric assemblies (21, 54, 67, 68). Thus, it is possible that VgrGi binds canonical VgrGs and prevents essential protein-protein interactions with other structural components of the T6SS. PAAR proteins, for instance, bind VgrG proteins at their C-termini (69). In *A. baylyi*, PAAR proteins are essential for effective T6SS firing (69). Thus, VgrG trimers containing VgrGi may be unable to bind PAAR proteins, which would be expected to disrupt T6SS assembly. Moreover, it is possible that VgrGi possesses conformational changes that abrogate critical baseplate-membrane complex interactions. Alternatively, it is possible that VgrGi acts post-T6SS assembly. The T6SS is expected to undergo two large conformational changes to enable deployment of the spiked tube. First, conformational changes in the baseplate are proposed to trigger sheath contraction (22, 70, 71). Secondly, the membrane complex is expected to undergo large conformational changes to enable the passage of the spiked tube (19, 72). It is possible that a leucine-to-arginine mutation causes aberrant interactions between VgrG and components of the membrane complex or baseplate, which prevent the necessary conformational changes for T6SS firing. In this instance, VgrGi may prove to be an invaluable tool to lock the dynamic T6SS in a static conformation and facilitate the structural characterization of the *Acinetobacter* T6SS.

Finally, it is noteworthy that we found an inactivating mutation of the T6SS in a clinical isolate. Although we cannot say with certainty whether this mutation was acquired during culturing or in the human host, it is tempting to speculate that there may be a selective pressure against the T6SS of medically relevant strains. As previously mentioned, the T6SS is an energetically expensive machine. Thus, there may be benefits to inactivating this machine, for example, to conserve energy. Moreover, T6SS structural components have been shown to be immunogenic (13, 73). It is possible that T6SS-inactive strains are selected for, as they are more likely to bypass detection by the immune system. In fact, most of the characterized virulence factors of *A. baumannii* involve evading immune detection or overcoming nutritional immunity (39). Thus, eliminating the T6SS may represent yet another strategy to bypass immune detection. Nonetheless, the T6SS confers a competitive advantage to environmental bacteria living in polymicrobial communities. Thus, it is likely that the environment exerts selective pressure in favor of a silenced T6SS in clinical *A. baumannii* strains and a constitutively active T6SS in non-pathogenic strains, as was previously proposed for *V. cholerae* (74). Indeed, clinical isolates of *A. baumannii* often have a non-functional T6SS due to genetic disruptions or absence of T6SS genes (45, 75–77). Our work indicates that point mutations at critical residues of T6SS structural proteins could constitute an underappreciated mechanism by which the T6SS is genetically disrupted.

Collectively, we have demonstrated that unlike in other bacteria, the C-terminus of VgrG is essential to functional T6SS assembly in *A. baumannii*. The specific roles of the DUF2345 and other C-terminal domains of VgrGs warrant further investigation.

## Materials and Methods

### Bacterial strains and growth conditions

All strains and plasmids used in this study are listed in Table S2. Strains were grown in Luria-Bertani (LB) broth at 37°C with shaking. Antibiotics were added to the media when appropriate (see below).

### Transposon mutagenesis and screen T6SS-active mutants

Our method for transposon mutagenesis has been described previously (42). Plasmid pSAM::OmpAp+Tn903 was introduced to AbCAN2 from *E. coli* BW19851 by bi-parental mating. Briefly, overnight cultures were pelleted, washed three times with fresh LB and resuspended at an OD600 of 1.0. The cultures were then mixed at a 1:1 ratio and spotted onto a dry LB-agar plate. Following a 5-h incubation at 37°C, the spot was resuspended in 1 ml of LB broth, and 10-fold dilutions were plated on LB-agar containing kanamycin (20 μg/ml) + chloramphenicol (12.5 μg/ml) and incubated overnight at 37°C. The transposon mutants obtained were subsequently subjected to a colony blot (52) to identify mutants with an active T6SS. Briefly, colonies were transferred to a nitrocellulose membrane and allowed to dry at room temperature for 30 minutes. The dried membrane is washed twice with TBST and treated as a normal Western blot, probing for Hcp and RNAP (see below). Candidates were then isolated and verified using a standard Hcp secretion assay (see below).

Identification of transposon insertion sites was carried out using a method adapted from (78). Transposon mutant gDNA was digested with MmeI and separated on a 0.7% agarose gel by electrophoresis. Fragments ∼500 and ∼1600 bp in length were extracted and appended with double-stranded oligonucleotide sequencing adapters by ligation. Finally, the transposon plus two 16-bp genomic sequences flanking the transposon was amplified by PCR and sequenced. The insertion site was verified by amplifying a genomic fragment of the putative insertion site by PCR and sequencing that fragment as well.

### Hcp secretion assay and Western blot

Overnight cultures were back-diluted in fresh LB to an OD600 of 0.025 and grown at 37°C with shaking until they reached an OD600 of 0.35-0.5 (AbCAN2) or 0.4-0.7 (Ab17978). Cells were then pelleted by centrifugation. Cells were resuspended in Laemmli buffer to a final OD of 10, while the supernatant fraction was centrifuged once again (as above) to pellet residual cells. Supernatant proteins were subsequently precipitated with trichloroacetic acid, as previously described (40). OD-normalized volumes of whole cells or supernatants were loaded onto 15% (for Hcp) or 8% (for VgrGs) SDS-PAGE gels for separation, transferred to a nitrocellulose membrane and probed with polyclonal rabbit anti-Hcp (1:1000, (40)), polyclonal rabbit anti-6xHis (1:2000, Invitrogen, Waltham, MA) or monoclonal mouse anti-RNA polymerase (1:2600, Biolegend, San Diego, CA). Western blots were then probed with IRDye-conjugated anti-mouse and anti-rabbit secondary antibodies (both at 1:15,000, LI-COR Biosciences, Lincoln, NE) and visualized with an Odyssey CLx imaging system (LI-COR Biosciences).

### Generation of mutants and pWH constructs

Primers used in this study are listed in Table S2. Marked mutant strains were generated by substitution of the gene of interest by a kanamycin resistance marker, as described previously (79). Selection was carried out using kanamycin (30 μg/ml). To generate clean mutants, electrocompetent marked mutants were transformed with pAT03 to remove the FRT-flanked kanamycin resistance cassette. Transformants were plated on LB-agar containing 2 mM IPTG + hygromycin (600 μg/ml). Mutant strains were verified by PCR and sequencing.

Constructs pWH-*vgrGi*-6xHis and pWH-*vgrG1*-6xHis were generated by restriction cloning using the EcoRI and PstI sites. These plasmids were then used as templates to make point mutants of VgrGi and VgrG1, respectively. Specifically, point mutants were generated using the QuikChange II Site-Directed Mutagenesis Kit (Agilent Technologies, Santa Clara, CA), according to the manufacturer’s instructions. Constructs of VgrGi truncations ΔDUF and ΔTT_R749L_ were generated by inverse PCR of pWH-*vgrGi*-6xHis followed by blunt ligation. All other VgrGi truncation constructs were generated by restriction cloning, as described above. Selection was carried out using tetracycline (15 μg/ml for AbCAN2 and 10 μg/ml for Ab17978). All constructs were verified by PCR and sequencing.

### Bacterial killing assay

Overnight cultures were washed three times with fresh LB and normalized to an OD600 of 1. Predator and prey strains were mixed at a 10:1 ratio, respectively, and spotted on dry LB-agar. Following a 3.5-h incubation at 37°C, spots were resuspended in LB and serial dilutions were spotted on kanamycin (30 μg/ml) LB-agar. CFUs of surviving prey cells were enumerated after overnight incubation at 37°C.

### VgrGi structural model and alignments

VgrGi C-gp5 (residues 1-666) was submitted to the iterative threading assembly refinement (I-TASSER) server (80), using the crystal structure of *P. aeruginosa* PAO1 VgrG1 (PDB: 4MTK) as a template without alignment. The trimer assembly was constructed by aligning the VgrGi model to each VgrG1^Pa^ monomer in Biological Assembly 1 (PDB: 4MTK), using the PyMOL Molecular Graphics System, Version 1.2r3pre (Schrödinger, LLC).

Domain designations were made according to Interpro (81) and HHPred (82) servers. The predictions for the coiled-coil domain and the DUF2345 domain overlap in residues 668-697; however, for the construction of the VgrGi truncations, preference was given to the DUF2345 domain. Sequence alignments were done using Clustal Omega (83).

## Acknowledgements

This work was supported by National Institutes of Health Grant 1R01AI125363-01 to M.F.F. J.L. is funded by the Washington University Chancellor’s Graduate Fellowship. The funders had no role in this study.

## Author contributions

All authors designed experiments and analyzed data; J.L. and P.M.L performed experiments; J.L. and M.F.F. wrote the manuscript.

---

**Table S1.** Comparison between T6SS-related genes of AbCAN2 and Ab17978.

| **Gene name<sup>#</sup>** | **Length (bp)** |  | **Identity** |
| --- | --- | --- | --- |
|  | **Ab17978/AbCAN2** | **Cover** |  |
| ACX60_11685 | 693 | 100 | 99 |
| TssB | 504 | 100 | 99 |
| TssC | 1482 | 100 | 99 |
| Hcp | 504 | 100 | 99 |
| TssE | 477 | 100 | 100 |
| TssF | 1812 | 100 | 99 |
| TssG | 999 | 100 | 99 |
| ACX60_11650 | 1413 | 100 | 98 |
| TssM | 3824 | 100 | 99 |
| TagF | 960 | 100 | 99 |
| TagN | 768 | 100 | 99 |
| PAAR | 264 | 100 | 99 |
| ClpV | 2682 | 100 | 98 |
| TssA | 1098/1095 | 99 | 99 |
| TssK | 1365 | 100 | 99 |
| TssL | 807 | 100 | 99 |
| ACX60_11605 | 603 | 100 | 99 |
| TagX | 954 | 100 | 99 |

<sup>#</sup> hypothetical proteins are named based on the locus tag of Ab17978

**Table S2.**
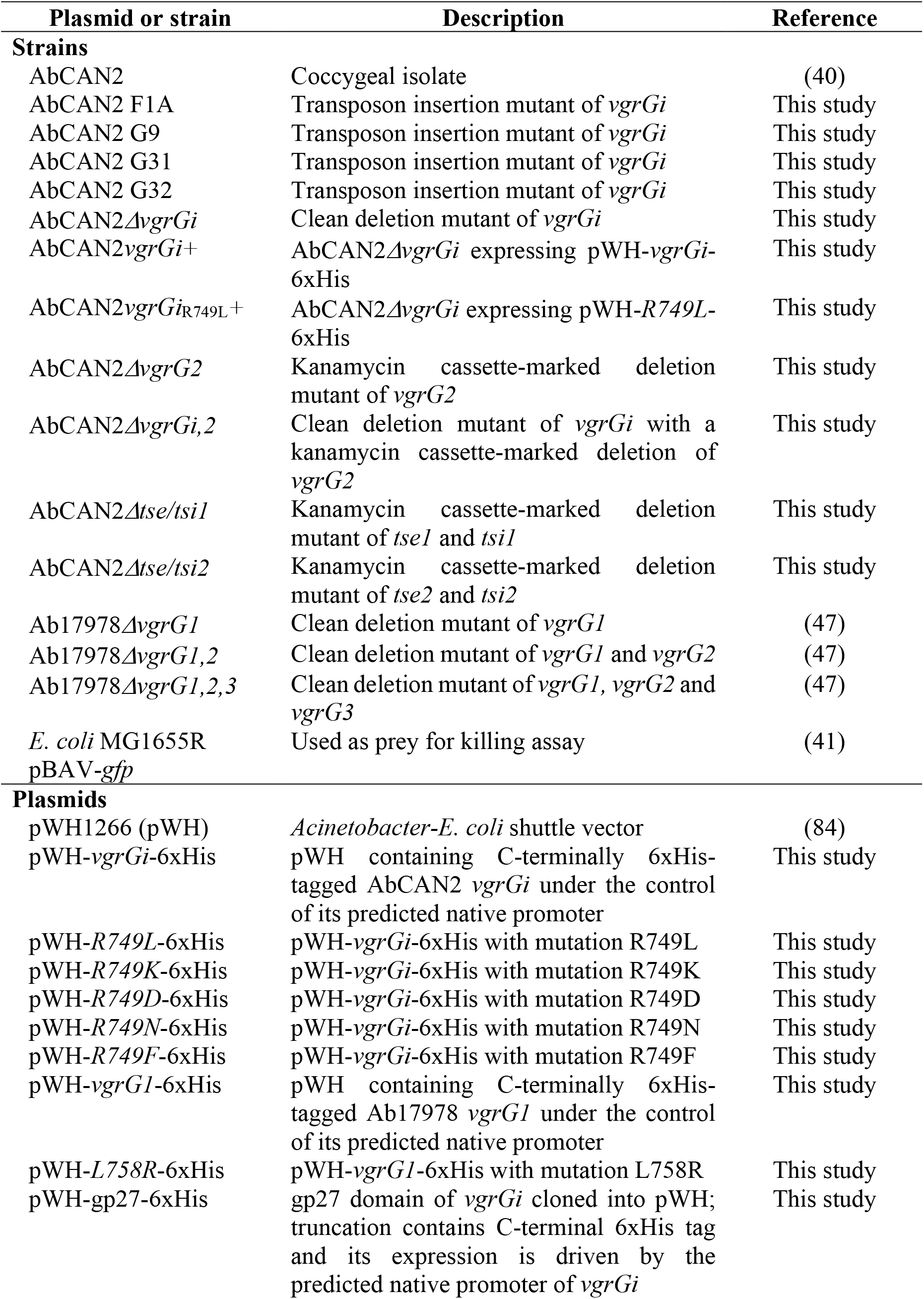
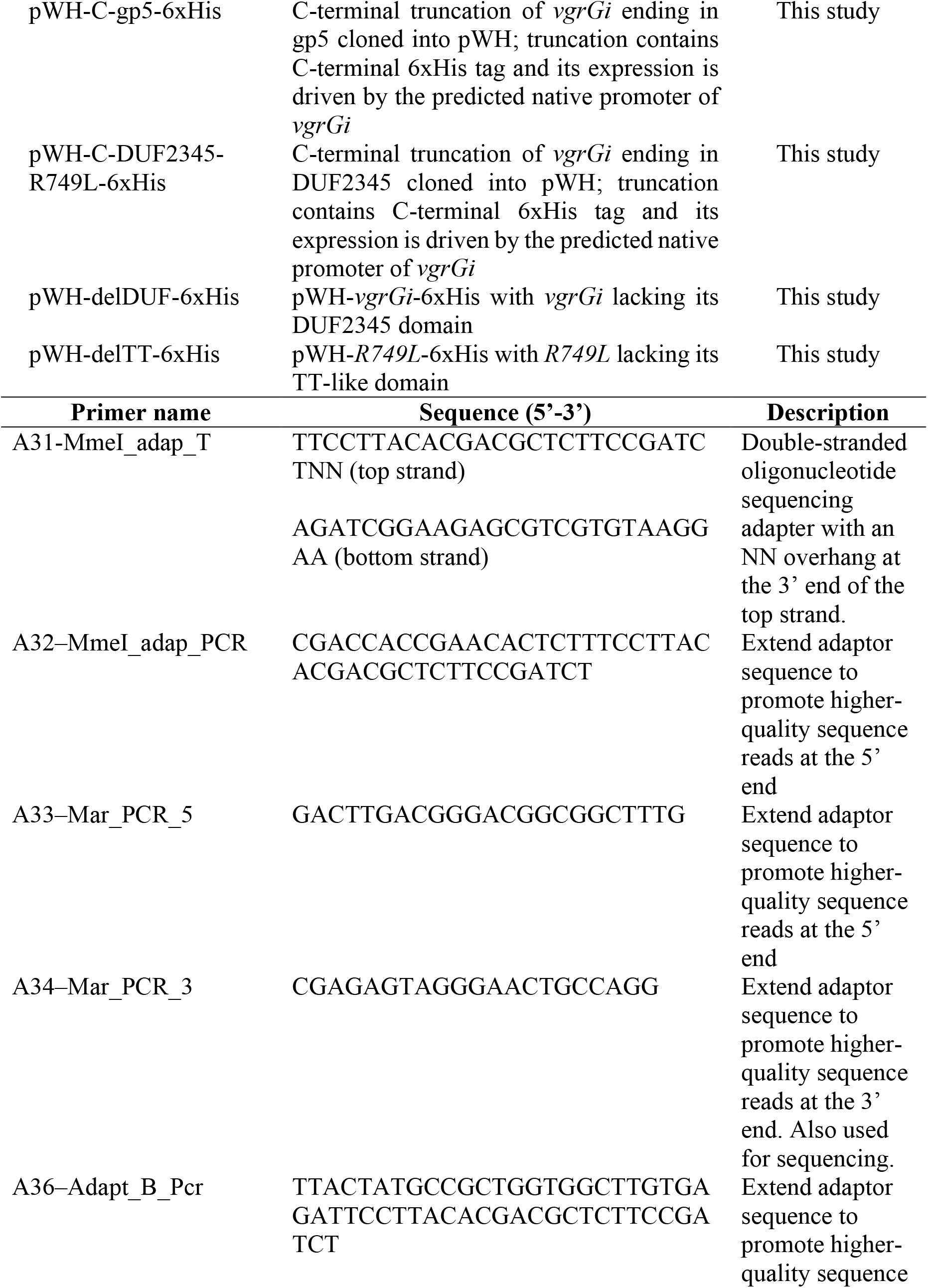
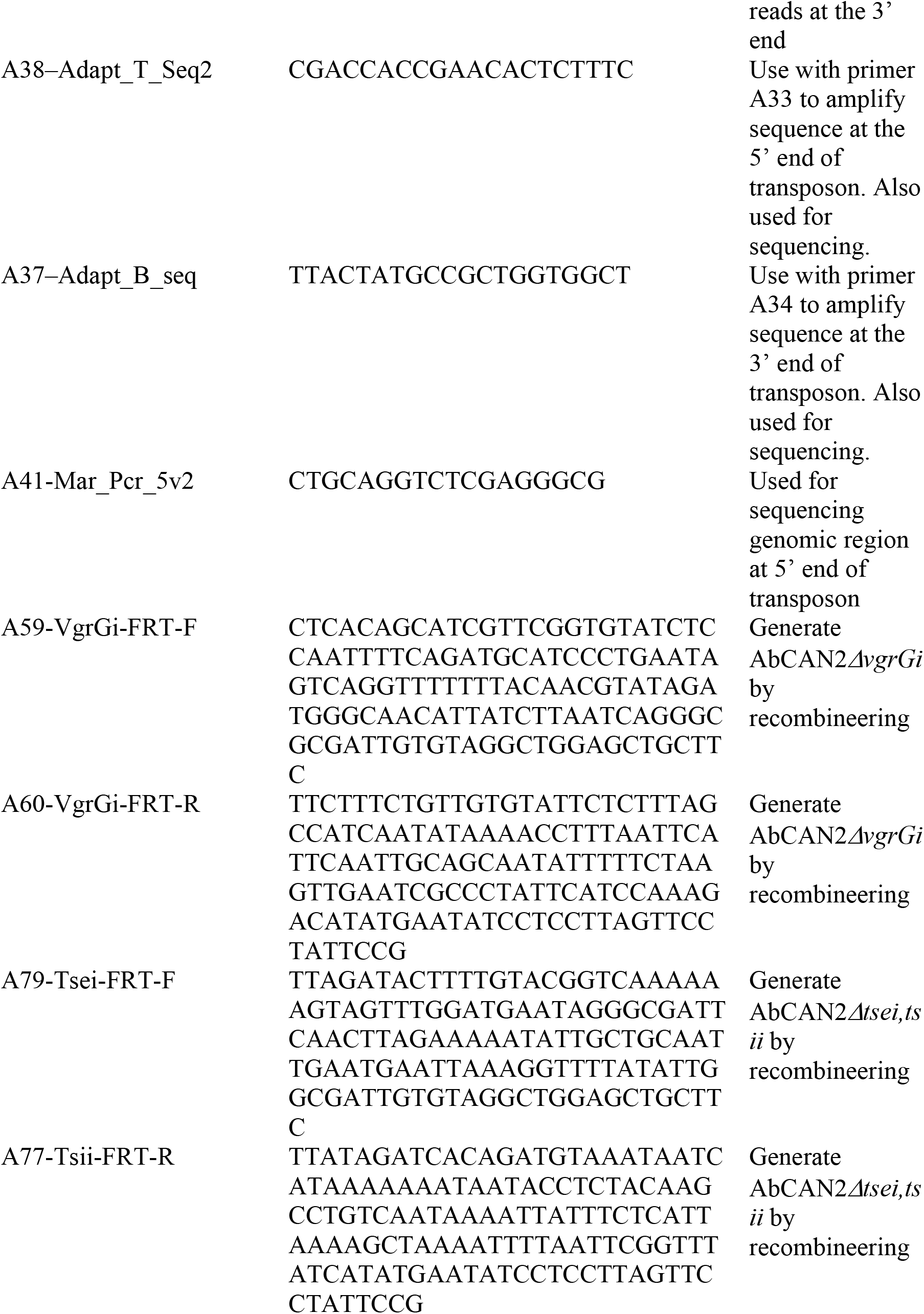
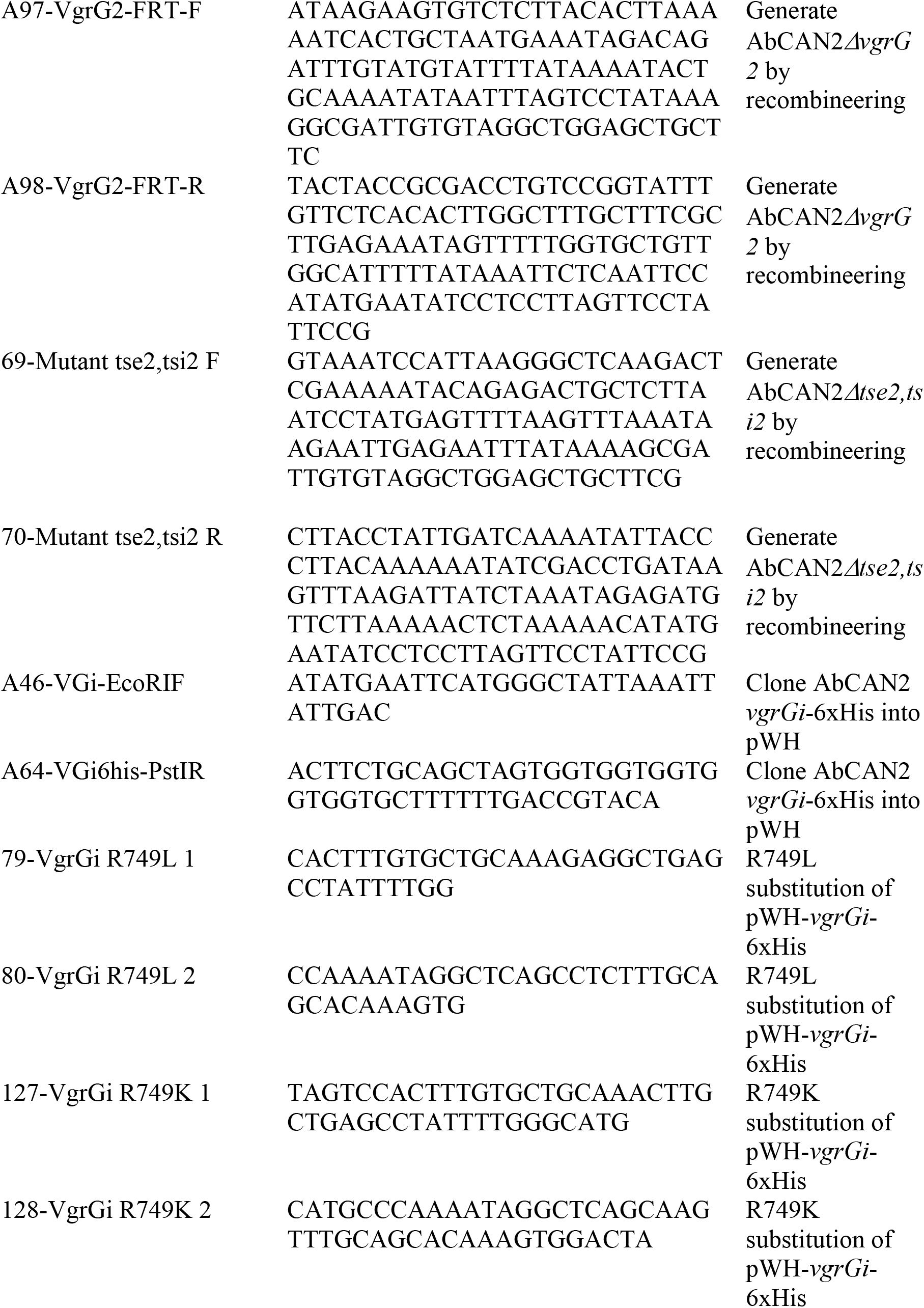
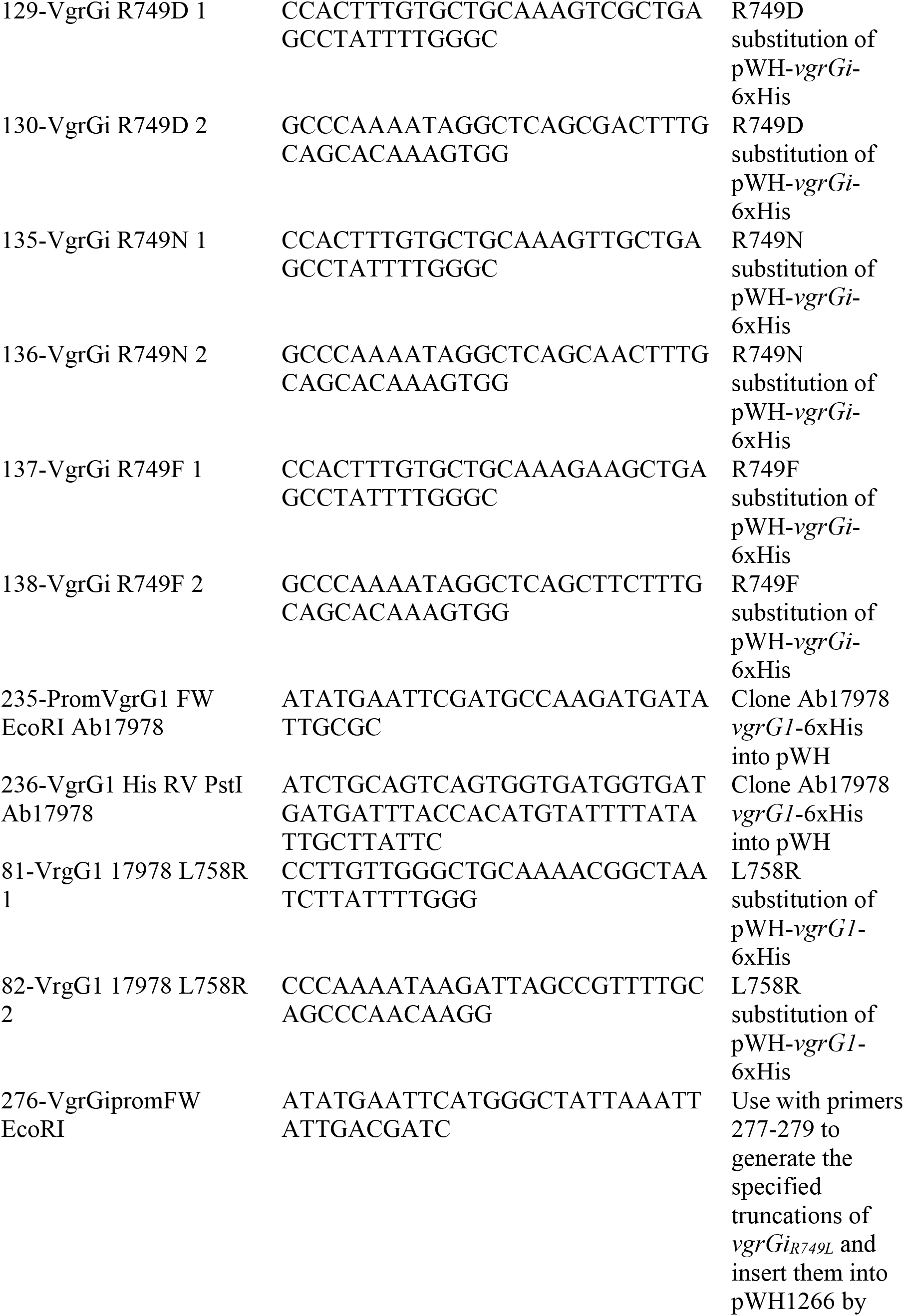
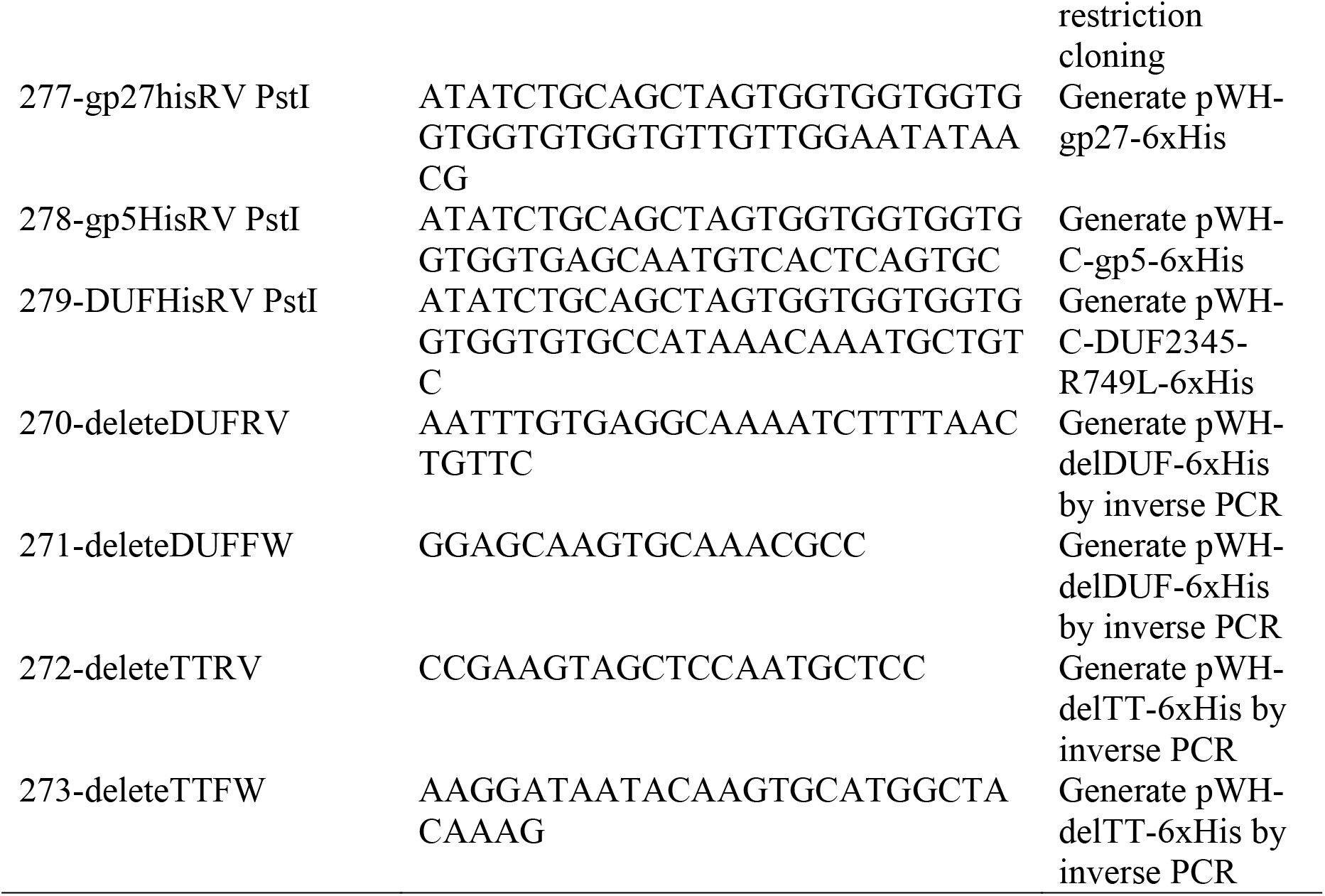
Strains, plasmids and primers used in this study.

**Figure S1.**
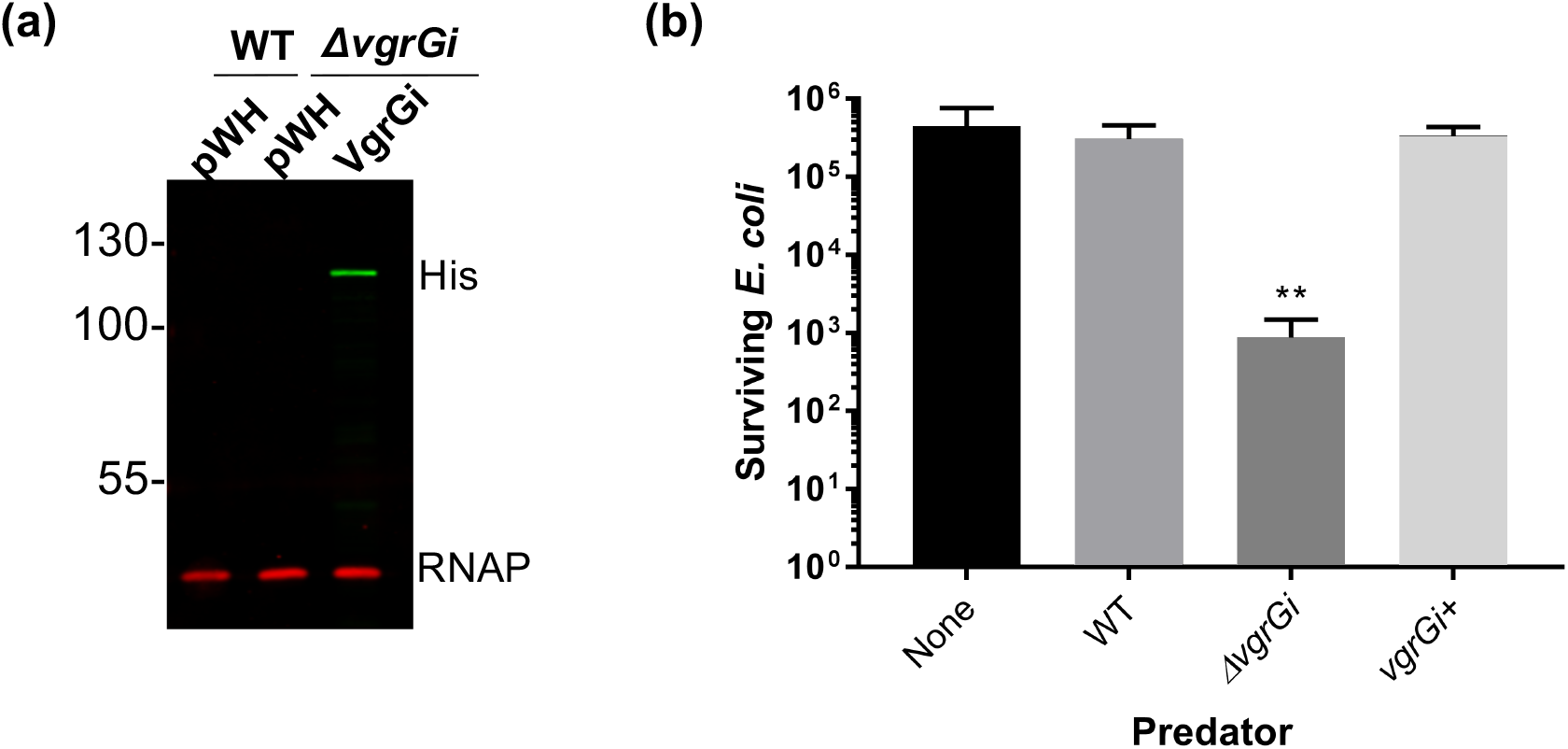
VgrGi inhibits the T6SS of AbCAN2. **a**, Western blot of OD-normalized whole cell fractions of the indicated strains probing for VgrGi-His expression. RNAP is included as a loading control. **b**, Cumulative data (mean ± SD) of 3 independent T6SS killing assays, each with technical duplicates. **, p = 0.0022 by Mann-Whitney test.

**Figure S2.**
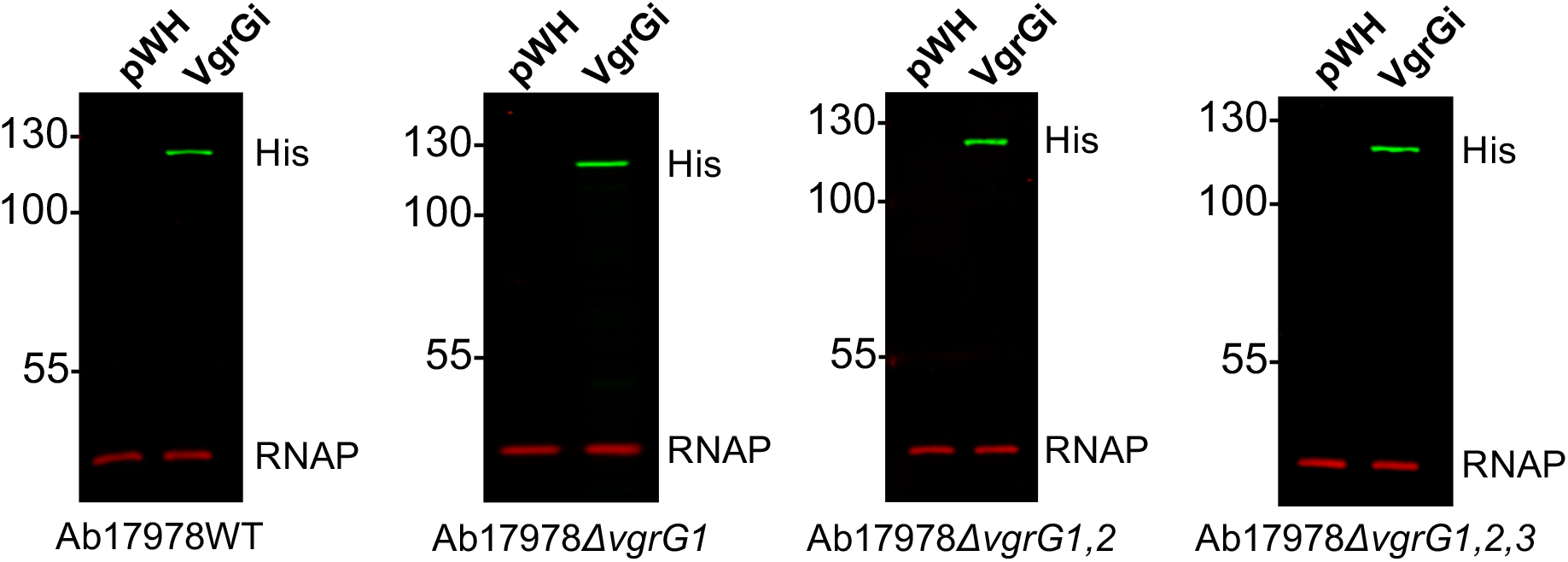
VgrGi expression in the indicated Ab17978 strains.

**Figure S3.**
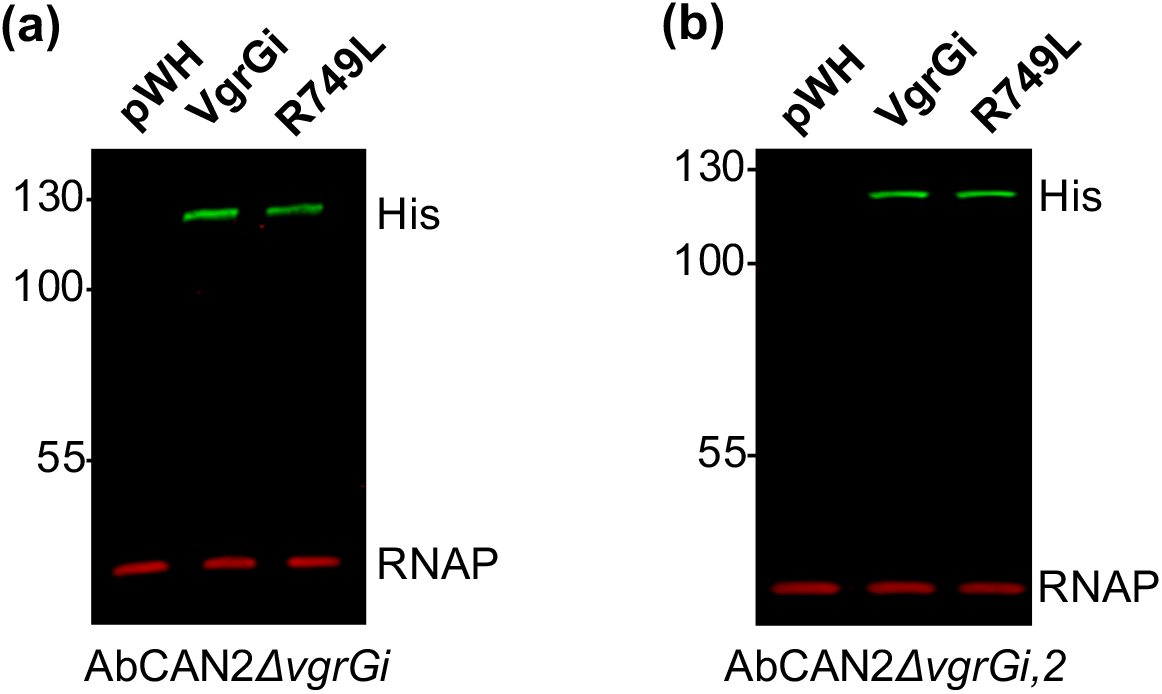
VgrGi/R749L expression in AbCAN2*ΔvgrGi* (a) or *ΔvgrGi,2* (b). pWH indicates a vector control. RNAP is included as a loading control.

**Figure S4.**
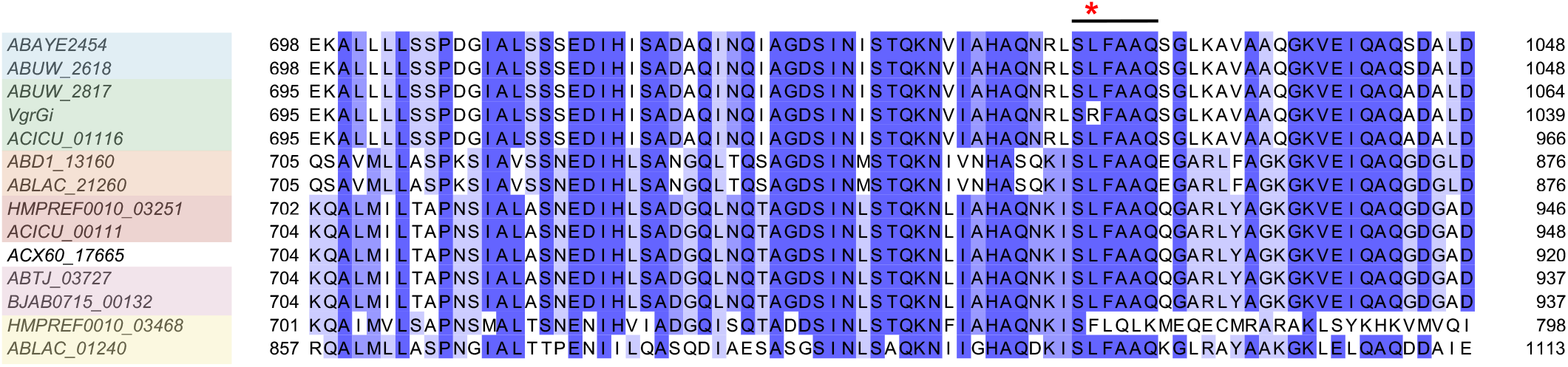
Amino acid alignment of VgrGi with VgrGs representating diverse phylogenetic groups. Residues are highlighted based on identity, with dark blue indicating residues that are highly conserved. The line indicates the SLFAAQ motif, while the asterisk indicates the position of R749 in VgrGi. VgrG locus tags are colored according to their phylogenic group (Fitzsimmons *et al.*, Infect Immun 86:e00297-18, 2018, https://doi.org/10.1128/IAI.00297-18): Class 1 (green), Class 2 (blue), Class 3 (yellow), Class 4 (light orange), Class 5 (pink) and Class 6 (dark orange). ACX60_17665 did not group with any of the other homologs.

**Figure S5.**
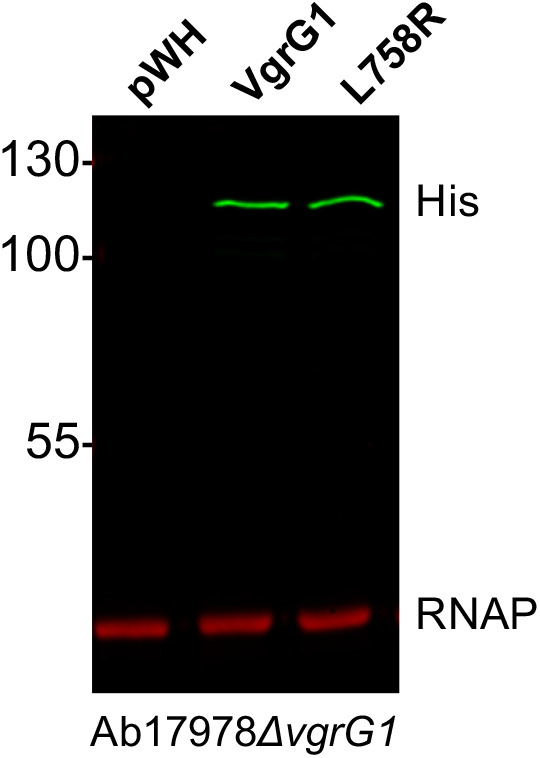
VgrG1/L758R expression in Ab17978*ΔvgrG1.* RNAP is included as a loading control.

**Figure S6.**
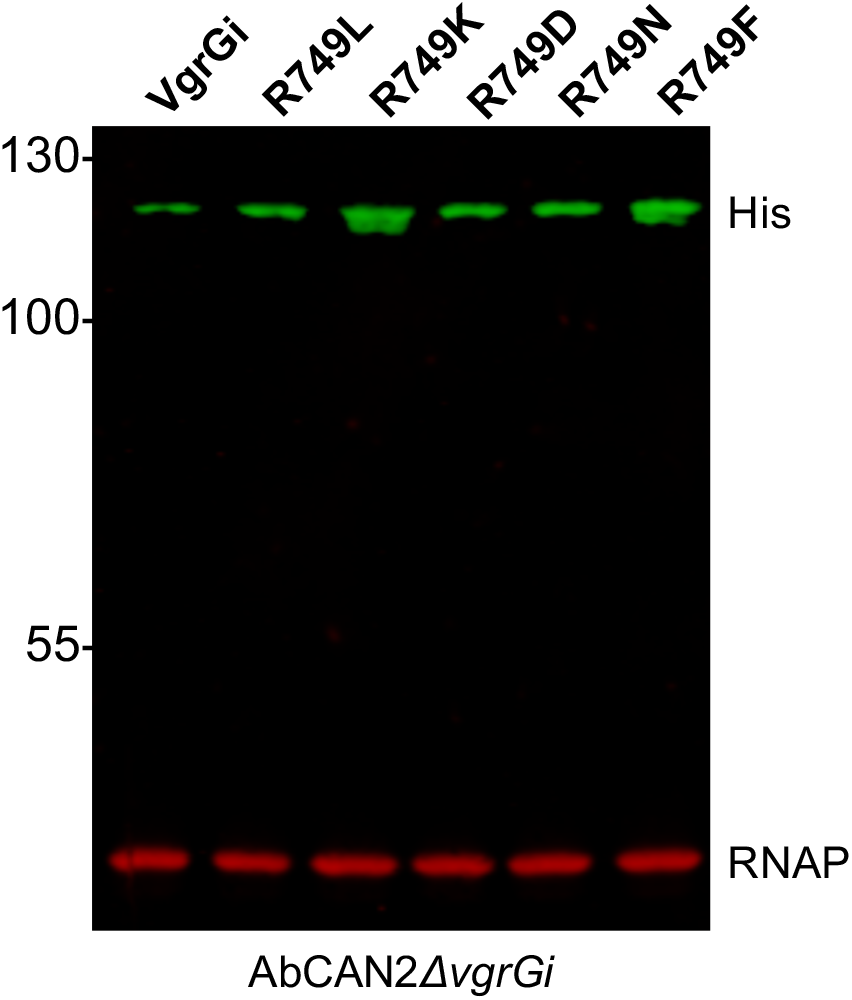
Expression of the indicated VgrGi point mutants in AbCAN2*ΔvgrGi*. RNAP is included as a loading control.

**Figure S7.**
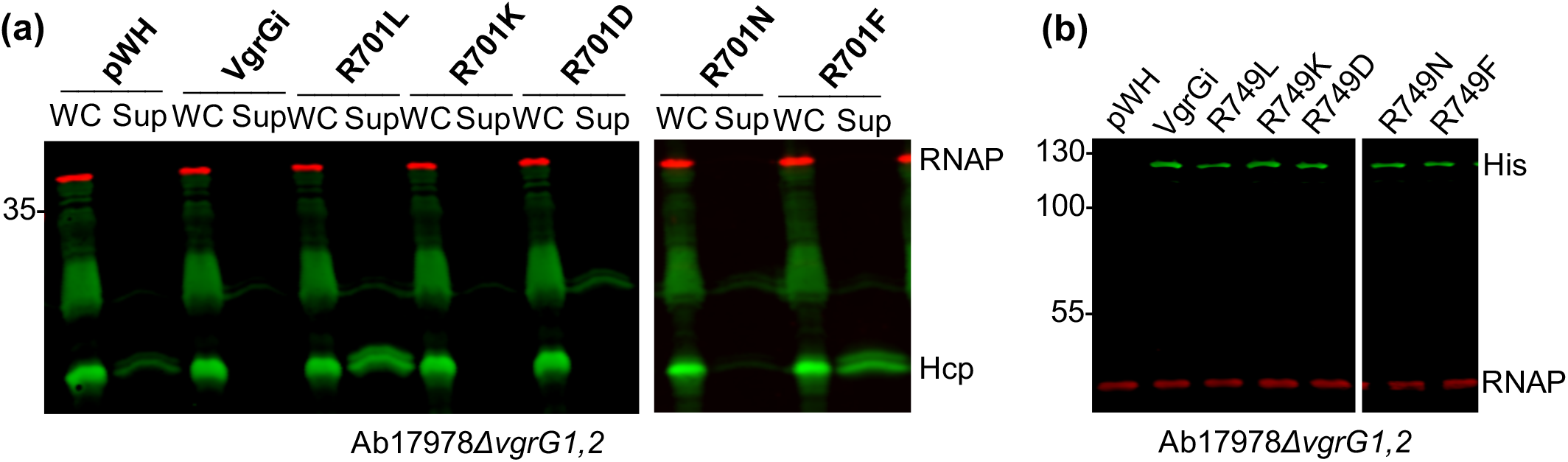
Polar and charged residues in position 749 result in T6SS inhibition by VgrGi. **a**, Western blot probing for Hcp expression and secretion. Shown are OD-normalized whole cell (WC) and supernatant (Sup) fractions of Ab17978*ΔvgrG1,2* expressing the indicated VgrGi point mutants. **b**, Expression of the indicated VgrGi point mutants in Ab17978*ΔvgrG1,2.* RNAP is included as a lysis and loading control.

**Figure S8.**
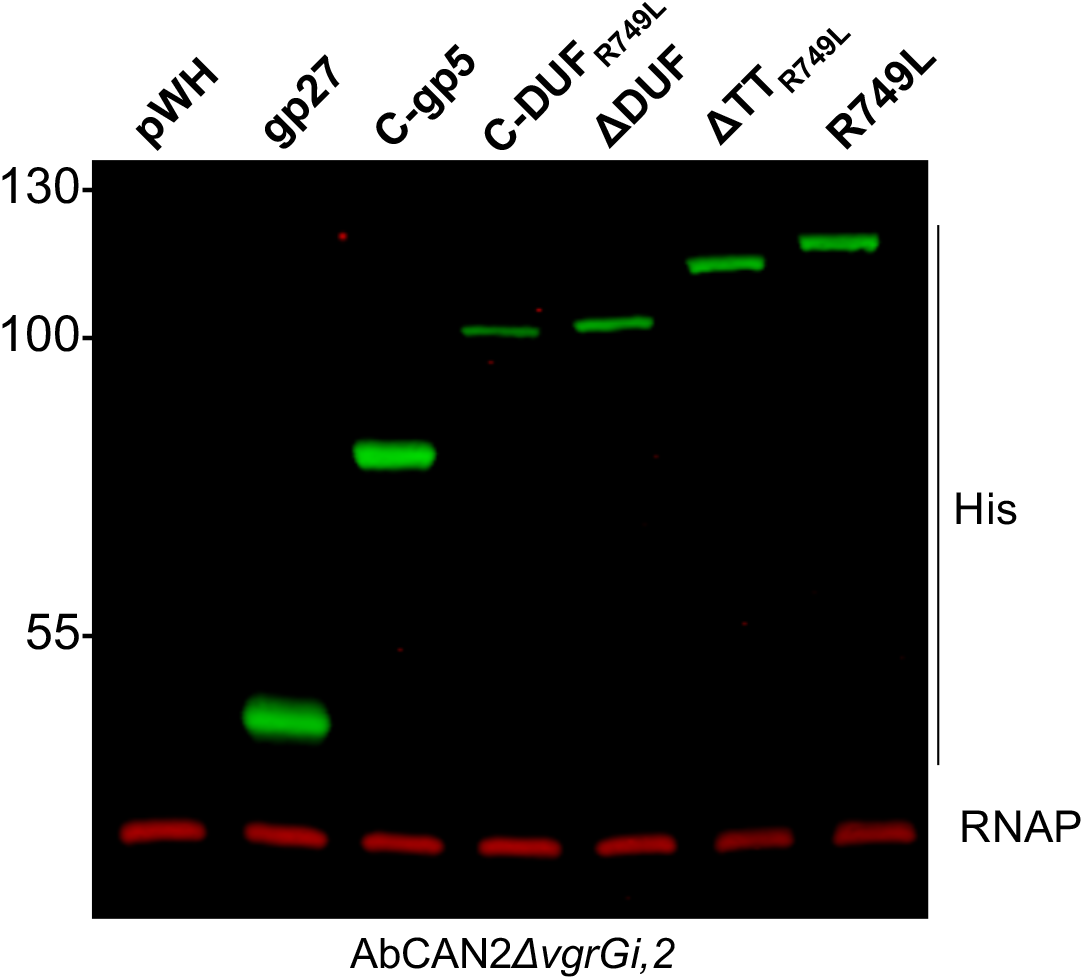
Expression of the indicated VgrGi_R749L_ truncations in AbCAN2*ΔvgrGi,2.* RNAP is included as a loading control.

## References

1. Hibbing ME, Fuqua C, Parsek MR, Peterson SB (2010) Bacterial competition: surviving and thriving in the microbial jungle. Nat Rev Microbiol 8(1):15–25.

2. Hood RD, Peterson SB, Mougous JD (2017) From Striking Out to Striking Gold: Discovering that Type VI Secretion Targets Bacteria. Cell Host Microbe 21(3):286–289.

3. Sana TG, et al. (2016) *Salmonella* Typhimurium utilizes a T6SS-mediated antibacterial weapon to establish in the host gut. Proc Natl Acad Sci 113(34):E5044–E5051.

4. Fu Y, Waldor MK, Mekalanos JJ (2013) Tn-seq analysis of vibrio cholerae intestinal colonization reveals a role for T6SS-mediated antibacterial activity in the host. Cell Host Microbe 14:652–663.

5. Anderson MC, Vonaesch P, Saffarian A, Marteyn BS, Sansonetti PJ (2017) Shigella sonnei Encodes a Functional T6SS Used for Interbacterial Competition and Niche Occupancy. Cell Host Microbe 21:769–776.

6. Borgeaud S, Metzger LC, Scrignari T, Blokesch M (2015) The type VI secretion system of Vibrio cholerae fosters horizontal gene transfer. Science (80-) 347(6217):63–67.

7. Thomas J, Watve SS, Ratcliff WC, Hammer BK (2017) Horizontal gene transfer of functional type VI killing genes by natural transformation. MBio 8(4):e00654–17.

8. Cooper RM, Tsimring L, Hasty J (2017) Inter-species population dynamics enhance microbial horizontal gene transfer and spread of antibiotic resistance. Elife 6:e25950.

9. Veening JW, Blokesch M (2017) Interbacterial predation as a strategy for DNA acquisition in naturally competent bacteria. Nat Rev Microbiol 15(10):621–629.

10. Hood RD, et al. (2010) A type VI secretion system of Pseudomonas aeruginosa targets a toxin to bacteria. Cell Host Microbe 7(1):25–37.

11. Jani AJ, Cotter PA (2010) Type VI Secretion: Not just for pathogenesis anymore. Cell Host Microbe 8(1):2–6.

12. Cianfanelli FR, Monlezun L, Coulthurst SJ (2016) Aim, Load, Fire: The Type VI Secretion System, a Bacterial Nanoweapon. Trends Microbiol 24(1):51–62.

13. Mougous JD, et al. (2006) A virulence locous of Pseudomonas aeruginosa encodes a protein secretion apparatus. Science (80-) 312:1526–1530.

14. Pukatzki S, et al. (2006) Identification of a conserved bacterial protein secretion system in Vibrio cholerae using the Dictyostelium host model system. Proc Natl Acad Sci 103(5):1528–1533.

15. Russell AB, Peterson SB, Mougous JD (2014) Type VI secretion system effectors: poisons with a purpose. Nat Rev Microbiol 12:137–148.

16. Durand E, Cambillau C, Cascales E, Journet L (2014) VgrG, Tae, Tle, and beyond: The versatile arsenal of Type VI secretion effectors. Trends Microbiol 22(9):498–507.

17. Basler M, Mekalanos JJ (2012) Type 6 Secretion Dynamics Within and Between Bacterial Cells. Science (80-) 337:815.

18. Brunet YR, Zoued A, Frederic B, Douzi B, Cascales E (2015) The type VI secretion TssEFGK-VgrG phage-like baseplate is recruited to the TssJLM membrane complez via multiple contacts and serves as assembly platform for tail tube/sheath polymerization. PLoS Genet 11(10):e1005545.

19. Durand E, et al. (2015) Biogenesis and structure of a type VI secretion membrane core complex. Nature 523:555–560.

20. Leiman PG, et al. (2009) Type VI secretion apparatus and phage tail-associated protein complexes share a common evolutionary origin. Proc Natl Acad Sci 106(11):4154–4159.

21. Pukatzki S, Ma AT, Revel AT, Sturtevant D, Mekalanos JJ (2007) Type VI secretion system translocates a phage tail spike-like protein into target cells where it cross-links actin. Proc Natl Acad Sci 104(39):15508–15513.

22. Cherrak Y, et al. (2018) Biogenesis and structure of a type VI secretion baseplate. Nat Microbiol:doi: 10.1038/s41564-018-0260-1. [Epub ahead of pri.

23. Nazarov S, et al. (2018) Cryo-EM reconstruction of Type VI secretion system baseplate and sheath distal end. EMBO J 37(4):1–14.

24. Zoued A, et al. (2016) Priming and polymerization of a bacterial contractile tail structure. Nature 531:59–63.

25. Basler M, Pilhofer M, Henderson GP, Jensen GJ, Mekalanos JJ (2012) Type VI secretion requires a dynamic contractile phage tail-like structure. Nature 483:182–186.

26. Basler M (2015) Type VI secretion system: Secretion by a contractile nanomachine. Philos Trans R Soc B Biol Sci 370:20150021.

27. Bernard CS, Brunet YR, Gueguen E, Cascales E (2010) Nooks and crannies in type VI secretion regulation. J Bacteriol 192(15):3850–3860.

28. Silverman JM, Brunet YR, Cascales E, Mougous JD (2012) Structure and regulation of the type VI secretion system. Annu Rev Microbiol 66:453–472.

29. Ishikawa T, et al. (2012) Pathoadaptive conditional regulation of the type VI secretion system in vibrio cholerae O1 strains. Infect Immun 80(2):575–584.

30. Miyata ST, Bachmann V, Pukatzki S (2013) Type VI secretion system regulation as a consequence of evolutionary pressure. J Med Microbiol 62:663–676.

31. Journet L, Cascales E (2016) The Type VI Secretion System in Escherichia coli and Related Species. EcoSal Plus 7(1):doi: 10.1128/ecosalplus.ESP-0009-2015.

32. Clemens DL, Lee B-Y, Horwitz MA (2018) The Francisella Type VI Secretion System. Front Cell Infect Microbiol 8(121). doi:10.3389/fcimb.2018.00121.

33. Joshi A, et al. (2017) Rules of Engagement: The Type VI Secretion System in Vibrio cholerae. Trends Microbiol 25(4):267–279.

34. Kostiuk B, Unterweger D, Provenzano D, Pukatzki S (2017) T6SS intraspecific competition orchestrates Vibrio cholerae genotypic diversity. Int Microbiol 20(3):130– 137.

35. Yang X, Pan J, Wang Y, Shen X (2018) Type VI Secretion Systems Present New Insights on Pathogenic Yersinia. Front Cell Infect Microbiol 8:260.

36. Brunet YR, Bernard CS, Gavioli M, Lloubès R, Cascales E (2011) An epigenetic switch involving overlapping fur and DNA methylation optimizes expression of a type VI secretion gene cluster. PLoS Genet 7(7):e1002205.

37. Ho BT, Basler M, Mekalanos JJ (2013) Type 6 secretion system-mediated immunity to type 4 secretion system-mediated horizontal gene transfer. Science (80-) 342(6155):250– 253.

38. Basler M, Ho BT, Mekalanos JJ (2013) Tit-for-tat: Type VI secretion system counterattack during bacterial cell-cell interactions. Cell 152(4):884–894.

39. Harding CM, Hennon SW, Feldman MF (2018) Uncovering the mechanisms of Acinetobacter baumannii virulence. Nat Rev Microbiol 16(2):91–102.

40. Weber BS, et al. (2013) Genomic and Functional Analysis of the Type VI Secretion System in Acinetobacter. PLoS One 8(1):e55142.

41. Weber BS, Ly PM, Irwin JN, Pukatzki S, Feldman MF (2015) A multidrug resistance plasmid contains the molecular switch for type VI secretion in *Acinetobacter baumannii*. Proc Natl Acad Sci 112(30):9442–9447.

42. Di Venanzio G, et al. (2019) Multidrug-resistant plasmids repress chromosomally encoded T6SS to enable their dissemination. Proc Natl Acad Sci 116(4):1378–1383.

43. Di Venanzio G, et al. (2019) Urinary tract colonization is enhanced by a plasmid that regulates uropathogenic Acinetobacter baumannii chromosomal genes. Nat Commun 10(2763). doi:10.1038/s41467-019-10706-y.

44. Repizo GD, et al. (2015) Differential role of the T6SS in Acinetobacter baumannii virulence. PLoS One 10(9):e0138265.

45. Kim J, et al. (2017) Microbiological features and clinical impact of the type VI secretion system (T6SS) in Acinetobacter baumannii isolates causing bacteremia. Virulence 8(7):1378–1389.

46. Hu Y, et al. (2018) Regulation of gene expression of hcp, a core gene of the type VI secretion system in Acinetobacter baumannii causing respiratory tract infection. J Med Microbiol 67:945–951.

47. Weber BS, et al. (2016) Genetic dissection of the type VI secretion system in Acinetobacter and identification of a novel peptidoglycan hydrolase, TagX, required for its biogenesis. MBio 7(5):e01253–16.

48. Ringel PD, Hu D, Basler M (2017) The Role of Type VI Secretion System Effectors in Target Cell Lysis and Subsequent Horizontal Gene Transfer. Cell Rep 21:3927–3940.

49. Tang JY, Bullen NP, Ahmad S, Whitney JC (2018) Diverse NADase effector families mediate interbacterial antagonism via the type VI secretion system. J Biol Chem 293(5):1504–1514.

50. Koskiniemi S, et al. (2013) Rhs proteins from diverse bacteria mediate intercellular competition. Proc Natl Acad Sci 110(17):7032–7037.

51. Alcoforado Diniz J, Coulthurst SJ (2015) Intraspecies competition in Serratia marcescens is mediated by type VI-secreted Rhs effectors and a conserved effector-associated accessory protein. J Bacteriol 197(14):2350–2360.

52. Weber BS, Ly PM, Feldman MF (2017) Screening for secretion of the type VI secretion system protein Hcp by enzyme-linked immunosorbent assay and colony blot. Bacterial Protein Secretion Systems: Methods and Protocols, pp 465–472.

53. Renault MG, et al. (2018) The gp27-like Hub of VgrG Serves as Adaptor to Promote Hcp Tube Assembly. J Mol Biol 430:3143–3156.

54. Wettstadt S, Wood TE, Fecht S, Filloux A (2019) Delivery of the Pseudomonas aeruginosa Phospholipase Effectors PldA and PldB in a VgrG- and H2-T6SS-Dependent Manner. Front Microbiol 10(July):1–18.

55. Flaugnatti N, et al. (2016) A phospholipase A1 antibacterial Type VI secretion effector interacts directly with the C-terminal domain of the VgrG spike protein for delivery. Mol Microbiol 99(6):1099–1118.

56. Brunet YR, Hénin J, Celia H, Cascales E (2014) Type VI secretion and bacteriophage tail tubes share a common assembly pathway. EMBO Rep 15(3):315–321.

57. Dong TG, Ho BT, Yoder-Himes DR, Mekalanos JJ (2013) Identification of T6SS-dependent effector and immunity proteins by Tn-seq in Vibrio cholerae. Proc Natl Acad Sci 110(7):2623–2628.

58. Spínola-Amilibia M, et al. (2016) The structure of VgrG1 from Pseudomonas aeruginosa, the needle tip of the bacterial type VI secretion system. Acta Crystallogr Sect D Struct Biol D72:22–33.

59. Fitzsimons TC, et al. (2018) Identification of Novel Acinetobacter baumannii Type VI Secretion System Anti-Bacterial Effector and Immunity Pairs. Infect Immun 86:e00297–18.

60. Russell AB, et al. (2014) A type VI secretion-related pathway in bacteroidetes mediates interbacterial antagonism. Cell Host Microbe 16(2):227–236.

61. Boyer F, Fichant G, Berthod J, Vandenbrouck Y, Attree I (2009) Dissecting the bacterial type VI secretion system by a genome wide in silico analysis: What can be learned from available microbial genomic resources? BMC Genomics 10. doi:10.1186/1471-2164-10-104.

62. Uchida K, Leiman PG, Arisaka F, Kanamaru S (2014) Structure and properties of the C-terminal β-helical domain of VgrG protein from Escherichia coli O157. J Biochem 155(3):173–182.

63. Schneider JP, et al. (2019) Diverse roles of TssA-like proteins in the assembly of bacterial type VI secretion systems. EMBO J e100825:1–17.

64. Ma L, Lin J, Lai E (2009) An IcmF Family Protein, ImpLM, Is an Integral Inner Membrane Protein Interacting with ImpKL, and Its Walker A Motif Is Required for Type VI Secretion System-Mediated Hcp Secretion in Agrobacterium tumefaciens. J Bacteriol 191(13):4316–4329.

65. Ma L, Narberhaus F, Lai E (2012) IcmF Family Protein TssM Exhibits ATPase Activity and Energizes Type VI Secretion. J Biol Chem 287(19):15610–15621.

66. Zheng J, Leung KY (2007) Dissection of a type VI secretion system in Edwardsiella tarda. 66(November):1192–1206.

67. Burkinshaw BJ, et al. (2018) A type VI secretion system effector delivery mechanism dependent on PAAR and a chaperone-co-chaperone complex. Nat Microbiol 3(5):632– 640.

68. Sana TG, et al. (2015) Internalization of Pseudomonas aeruginosa Strain PAO1 into Epithelial Cells Is Promoted by Interaction of a T6SS Effector with the Microtubule Network. MBio 6(3):e00712–15.

69. Shneider MM, et al. (2013) PAAR-repeat proteins sharpen and diversify the type VI secretion system spike. Nature 500(7462):350–353.

70. Leiman PG, et al. (2010) Morphogenesis of the T4 tail and tail fibers. Virol J 7:1–28.

71. Taylor NMI, et al. (2016) Structure of the T4 baseplate and its function in triggering sheath contraction. Nature 533:346–352.

72. Rapisarda C, et al. (2019) In situ and high-resolution cryo-EM structure of a bacterial type VI secretion system membrane complex. EMBO J 38(10):1–18.

73. Zheng J, Shin OS, Cameron DE, Mekalanos JJ (2010) Quorum sensing and a global regulator TsrA control expression of type VI secretion and virulence in Vibrio cholerae. Proc Natl Acad Sci U S A 107(49):21128–21133.

74. Miyata ST, Bachmann V, Pukatzki S (2013) Type VI secretion system regulation as a consequence of evolutionary pressure. J Med Microbiol 62:663–676.

75. Wright MS, et al. (2014) New insights into dissemination and variation of the health care-associated pathogen Acinetobacter baumannii from genomic analysis. MBio 5(1):1–14.

76. Meumann EM, et al. (2019) Genomic epidemiology of severe community-onset Acinetobacter baumannii infection. Microb genomics 5(3):1–13.

77. Traglia G, et al. (2018) Genome sequence analysis of an extensively drug-resistant Acinetobacter baumannii indigo-pigmented strain depicts evidence of increase genome plasticity. Sci Rep 8(16961). doi:10.1038/s41598-018-35377-5.

78. Goodman AL, et al. (2009) Identifying Genetic Determinants Needed to Establish a Human Gut Symbiont in Its Habitat. Cell Host Microbe 6:279–289.

79. Tucker AT, et al. (2014) Defining gene-phenotype relationships in Acinetobacter baumannii through one-step chromosomal gene inactivation. MBio 5(4):e01313–14.

80. Roy A, Kucukural A, Zhang Y (2010) I-TASSER: A unified platform for automated protein structure and function prediction. Nat Protoc 5(4):725–738.

81. Mitchell AL, et al. (2019) InterPro in 2019: Improving coverage, classification and access to protein sequence annotations. Nucleic Acids Res 47:D351–D360.

82. Söding J, Biegert A, Lupas AN (2005) The HHpred interactive server for protein homology detection and structure prediction. Nucleic Acids Res 33:W244–W248.

83. Sievers F, et al. (2011) Fast, scalable generation of high-quality protein multiple sequence alignments using Clustal Omega. Mol Syst Biol 7(539). doi:10.1038/msb.2011.75.

84. Hunger M, Schmucker R, Kishan V, Hillen W (1990) Analysis and nucleotide sequence of an origin of an origin of DNA replication in Acinetobacter calcoaceticus and its use for Escherichia coli shuttle plasmids. Gene 87:45–51.

